# Visual Landmarks Influence Eye-Hand Coordination in Memory-Guided Reaches

**DOI:** 10.64898/2026.11.13.766551

**Authors:** Jennifer Y. X. Lin, Verónica Nácher, Ashkan Karimi, Hongying Wang, Saihong Sun, Xiaogang Yan, J. Douglas Crawford

## Abstract

The human visual system optimally weighs egocentric cues (e.g., object location relative to the eye) and allocentric cues (e.g., object location relative to other landmarks) to aim actions. The neural mechanisms of this process have been studied in the primate gaze system, but it is not known how these results extend to eye-hand coordination and the reach system. Here, reach and gaze (eye-in-space) data were collected from two head-unrestrained *Macaca mulatta* (M1, M2) trained to reach toward remembered visual targets on a touch screen. During the delay, animals either saw nothing (no-landmark), a pattern of ‘dots’ around the target (stable landmark), or a similar landmark that reappeared at a shifted position (landmark shift). In most conditions, increased delay times (up to 1200 ms) reduced reaction times, with minimal effects on gaze and reach errors. In the temporal domain, landmarks reduced gaze reaction time, increased the time to final hand touch, and reduced gaze-hand coupling. In the spatial domain, the stable landmark tended to improve reach accuracy and precision but reduced landmark-centered gaze-hand coupling. Landmark shifts increased errors in pooled datasets, due to weighted (and modestly correlated) shifts in both reach and gaze in the same direction. The ‘allocentric weights’ for reach were also modulated by initial gaze position. Overall, these data suggest that visual landmarks influence the timing, accuracy, and coordination of gaze / hand movements, likely through visual feedback during reach, and these effectors share a similar mechanism for optimal integration of egocentric and allocentric visual cues.

**NEW & NOTEWORTHY STATEMENT:** Visual landmarks influence action goals, but their influence on eye-hand coordination has not been studied. Here, when monkeys reached toward remembered targets, stable landmarks reduced gaze reaction times and spatiotemporal coupling of the eye and hand and tended to improve reach accuracy and precision. Landmark shifts induced correlated gaze and reach endpoint shifts in the same direction. These results replicate landmark influences on human reach goals, extend this to eye-hand coordination, and establish a potential experimental model for neurophysiological investigation.

## INTRODUCTION

The visual system has access to two types of spatial information: egocentric, i.e., object location relative to the eye, and allocentric, i.e., object location relative to other visual stimuli, i.e., landmarks (Ruotolo et al., 2011; Simons & Wang, 1998). Whereas allocentric codes are associated with the ventral (occipital-temporal) visual stream, egocentric processing is associated with the dorsal visual stream (Goodale & Milner, 1992; Schenk, 2006). The use and integration of these spatial cues for perception, memory, and navigation systems is well documented (Bird & Burgess, 2008; Burgess, 2006; Byrne et al., 2007; McNaughton et al., 2006) but for many years the sensorimotor community focused almost completely on egocentric representations (Andersen & Buneo, 2002; Buneo & Andersen, 2006; Crawford et al., 2011). More recent studies have shown that egocentric and allocentric cues are integrated to provide optimal reach behavior in the human (Byrne & Crawford, 2010; Fiehler et al., 2014). This has inspired neuroimaging studies in humans (Chen et al., 2014; Musa et al., 2025) and neurophysiological mechanisms in monkeys (Bharmauria et al., 2025), but so far, the neurophysiology of ego / allocentric integration for eye-hand coordination and reaches is poorly understood.

Human studies have shown that the presence of stable allocentric cues benefits the accuracy, precision, and timing of reaches, especially in the presence of a memory delay (Byrne & Crawford, 2010; Chen et al., 2011; Fiehler et al., 2014; Klinghammer et al., 2015, 2017; Krigolson & Heath, 2004; Lu & Fiehler, 2020; Musa et al., 2024; Obhi & Goodale, 2005). Byrne & Crawford (2010) showed that this occurs through an optimal integration process: when ego / allocentric cues conflict, the brain weights each according to their relative reliability and perceived stability. Subsequent cue-conflict studies employed virtual reality to show that this weighting is also influenced by other factors such as proximity, depth, and even scene semantics (Baltaretu et al., 2024; Fiehler et al., 2014; Klinghammer et al., 2016, 2017; Lu & Fiehler, 2020). Further neuroimaging experiments suggest that this process involves the convergence of ventral stream allocentric signals with dorsal stream egocentric signals within parietal and frontal cortex (Chen et al., 2018; Musa et al., 2025).

The primate visuomotor system has been used to investigate complex stimulus interactions at the cellular level, including object-centered codes for saccade targets (Olson & Gettner, 1999) and reach targets (Taghizadeh et al., 2024). A behavioral study in monkeys found that visual landmarks also stabilize gaze aiming (Li et al., 2017), and used the cue-conflict approach to show that the gaze system uses a similar weighting process to that observed in human reaches, modulated by spatial relationships between initial gaze, target, and landmark position. This enabled a series of neurophysiological experiments in the frontal and supplementary eye fields, showing that 1) target and landmark signals persist and interact to provide landmark-centered codes in prefrontal visual responses (A. Schütz et al., 2023), 2) delay responses encode ego / allocentric cue conflicts, and 3) these conflicting codes are re-integrated at the cellular level in the gaze motor response (Bharmauria et al., 2020, 2021). However, to our knowledge, landmark-centered coding has not been investigated in the primate reach system.

To investigate the cellular basis of a sensorimotor transformation, it is important to first have a thorough understanding of the model system’s behavioral output. In this case, it would be important to understand 1) if landmarks somehow benefit reach behavior in monkeys, 2) if landmarks are weighted with egocentric cues in the cue-conflict paradigm, 3) how these interactions are influenced by different memory delays, and 4) how these interactions impact the complex spatiotemporal relationships between gaze and reach behavior (Arora et al., 2019; Crawford et al., 2004; Johansson et al., 2001), including the use of gaze signals to guide the hand (Henriques & Crawford, 2000; Neggers & Bekkering, 2000; Prablanc et al., 1979). Based on the notion that the gaze and reach systems share high level visual computations but more independent motor mechanisms (Crawford et al., 2004; de Brouwer et al., 2021; Snyder et al., 2002), one might expect to observe similar ego / allocentric interactions to those found in humans and the primate gaze system, but with minimal effect on the motor aspects of eye-hand coordination.

The current study addressed this knowledge gap by recording gaze and reach data from macaque monkeys engaged in a memory guided reach task in the dark or in the presence of landmarks that could be stable or shift to produce cue conflict. Animals’ heads were unrestrained, so we focused on gaze (eye in space) and hand movement. Based on the literature reviews above, we tested (1) if a stable landmark decreased variable error, constant error, and reaction times, (2) how reach timing and position correlate with gaze in space or landmark-centered coordinates, (3) if normal gaze-hand coordination strategies would be preserved in the presence of landmarks, (4) if gaze and reaches deviate the same way during landmark shifts, and (5) how these deviations depend on the spatial details of target position, landmark position, and landmark shift direction. In general, most of these predictions were borne out, along with several unexpected observations, in particular an influence of landmarks on the spatiotemporal coupling of gaze and hand movements.

## METHODS

### Animals and Surgical Procedures

Data were collected from two female rhesus macaque monkeys (*Macaca mulatta*), henceforth referred to as M1 and M2. The animals weighed around 7 kg at the time of the study. The animals underwent several surgeries before the experiment, some described previously (Arora et al., 2019). Under general anesthesia (1.5% isoflurane following intramuscular injection of 10 mg/kg ketamine hydrochloride, 0.05 mg/kg atropine sulfate, and 0.5 mg/kg acepromazine) a steel head post was implanted onto the skull using an acrylic head cap secured by screws. One Teflon-coated search coil was implanted in the eye subconjunctivally to record horizontal and vertical 2D eye orientation. The animals then rested for two weeks after surgery to recover. During data collection, fluid intake was controlled as the animal received fluid as a reward for correct trials. All of the surgical and experimental protocols were done according to the Canadian Council for Animal Care guidelines. The use of laboratory animals was reviewed and approved by the York University animal care committee.

### Experimental Setup and Behavioural Recordings

The experimental setup was similar to the one used by Arora et al. (2019) (Figure 1). During the experiment, the animal was seated in a customized Crist primate chair that allows for free motion of the head and forward reaching movements. The animal wore a removable head restraint, removable top plates, a security vest, and a shoulder collar.

**Figure 1.**
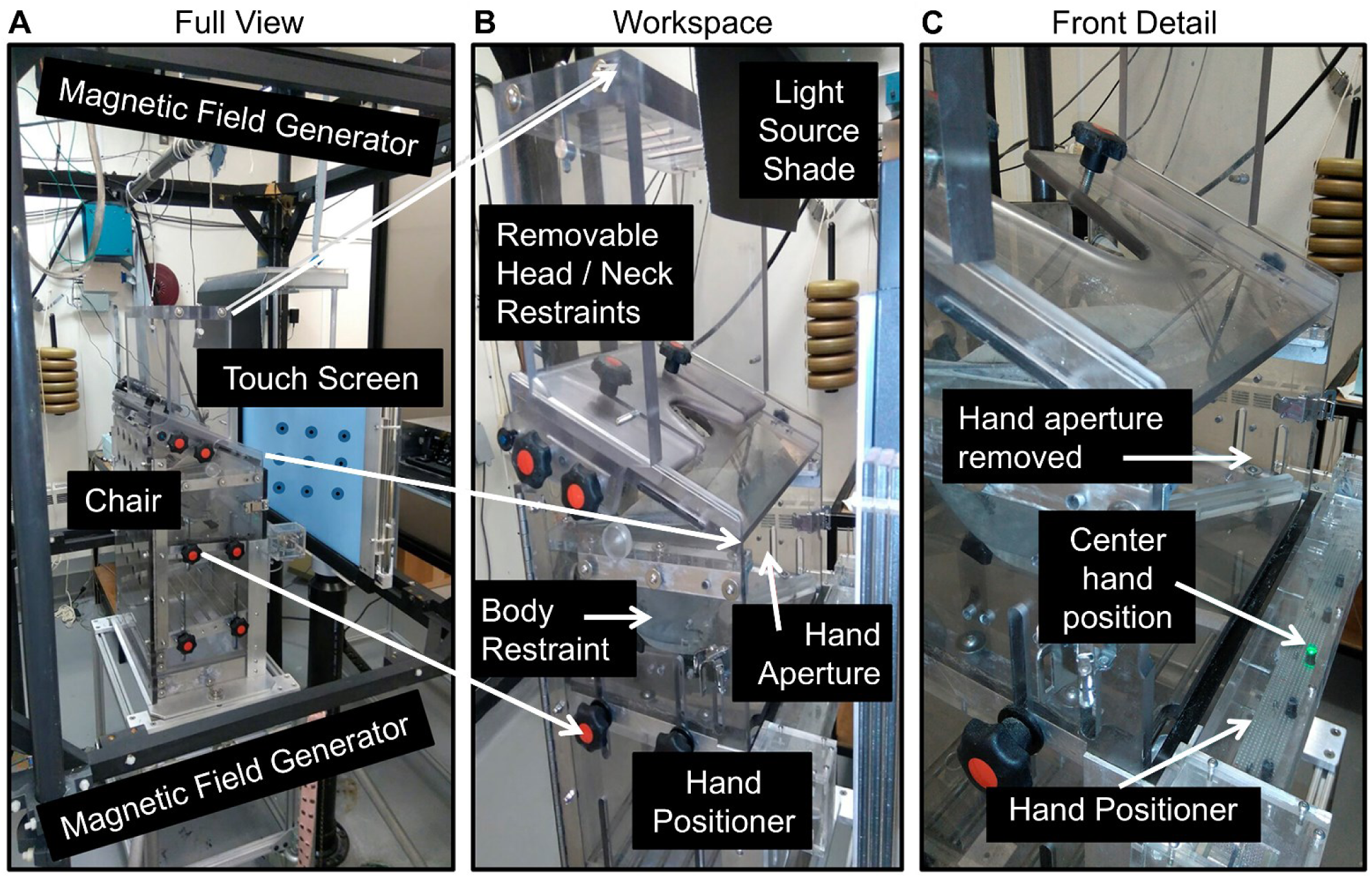
Laboratory setup. (A) A touch screen was placed in front of the customized Crist primate chair. (B) The removable head/neck restraints were removed during the experiment to allow for movement of the head. The body restraint was a jacket attached to the inside of the primate chair. This secured the monkey’s body position. Removing the hand aperture allowed movement of the hand. (C) The hand positioner was composed of the LED bar positioned at waist level and in front of the hands. The green LED light indicated the center hand position. Adapted from “Eye-head-hand coordination during visually guided reaches in head-unrestrained macaques”, by Arora, H. K., Bharmauria, V., Yan, X., Sun, S., Wang, H., & Crawford, J. D., 2019, *Journal of Neurophysiology*, *122*(5), 1946–1961. Copyright 2019 by the American Physiological Society.

Additional restrictions included a chain that attached the animal’s vest to the chair and an upper arm Velcro cuff connected to the chair that restrains the left arm.

The animal was trained to rest its hand on an LED bar attached to the chair at waist level. The center LED was 16 cm ahead and 28 cm below the right shoulder. The three LEDs we used are at (-19 °, -64 °), (19 °, -64 °), and (45 °, -64 °) in horizontal / vertical visual angles, respectively. We used a pair of infrared emitter and sensor to monitor if the hand was resting on the LED bar properly (in the vertical dimension), and a light sensor array above the LED bar to detect whether the animals’ hand is located on the initial position indicator LED (in horizontal dimension). Visual stimuli were displayed using a laser projector that projected onto a screen that was 27 cm distance ahead of the eye. An IR touch screen frame, with horizontal and vertical sensors, was attached in front of the screen to measure the 2D location of the hand contact on the screen.

The steel head post was used to stabilize the animal’s head during setup. The head coil, 2D eye coil, reward tube, and fluid dispenser were attached to the head cap. Two calibrations were done before the experiment. The 2D magnetic fields were precalibrated by rotating an external coil through each field direction and adjusting the field strength in each direction to an equal maximum (Arora et al., 2019). The second calibration required the animal to fixate on each of the nine targets displayed on the screen for 1 s.

### Behavioural Paradigm

The experiment was conducted in complete darkness to eliminate other stimuli. All visual stimuli were presented in red, with the target stimulus having the highest intensity. The egocentric cue was a dot at the center of the screen where the monkey would fixate their gaze during each trial (Figure 2A). The allocentric cue, known as the landmark, was four identical dots positioned at the vertices of a square with a 10 ° edge length. The landmark was located at 15 different locations within a 50 ° by 30 ° rectangle (Figure 2A). The target was a dot located at a random location within a 5 ° diameter circle at the center of the landmark. We also included “catch trials” with targets outside of the landmark to ensure animals reached towards the target, rather than the center of the landmark (Figure 2B).

**Figure 2.**
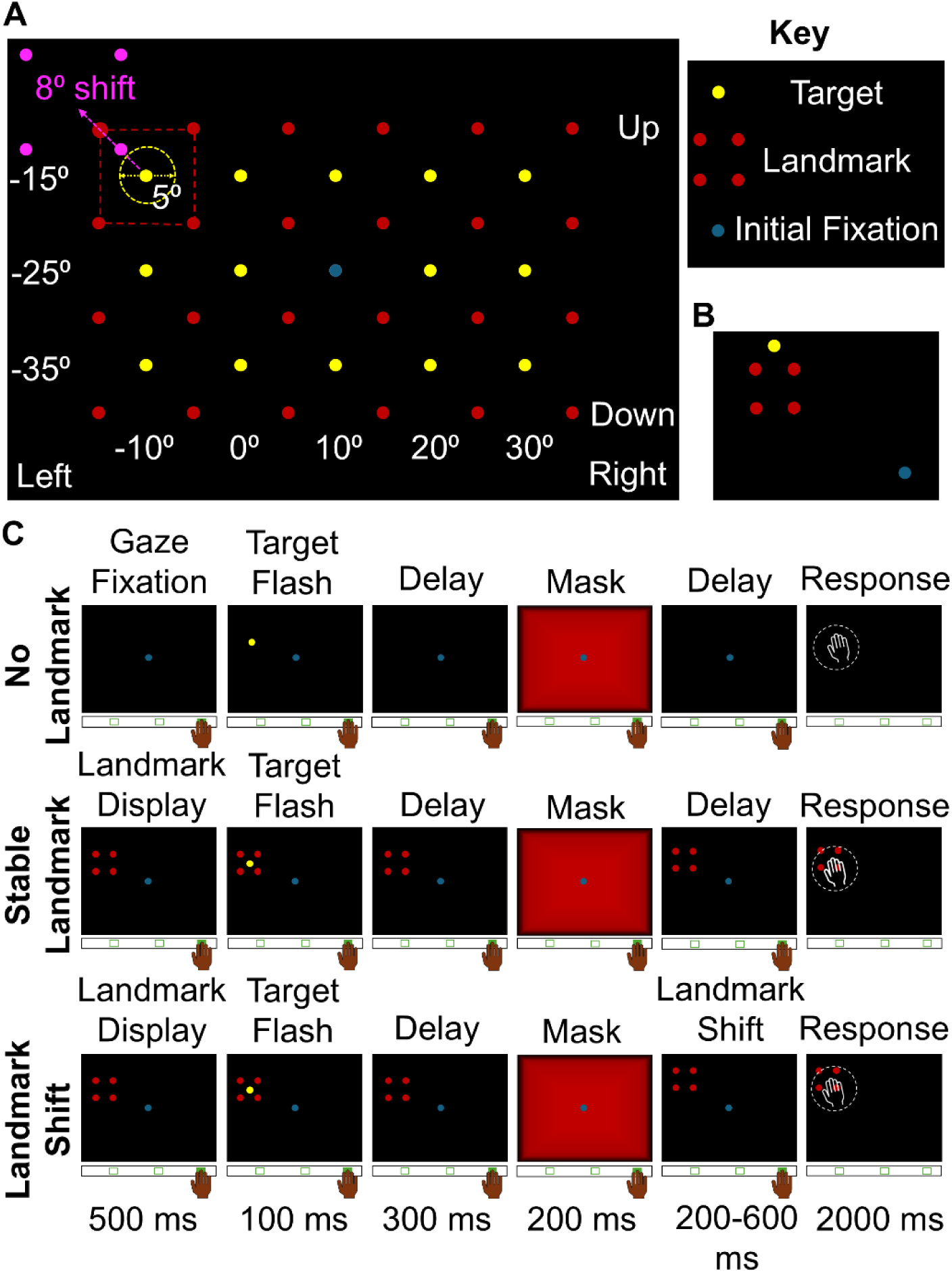
Visual Stimuli and Experimental Paradigm. (A) Possible locations of the target and landmark. This figure shows the fifteen locations where the target and landmark appeared during each trial. The corresponding target was located within a 5 ° diameter circle at the center of the landmark. (B) An example of a “catch trial” where the target is located outside of the landmark. (C) The experimental paradigm shows the time course and visual information shown to the monkey during the no-landmark condition, stable landmark condition, and landmark shift condition. The blue dot is the gaze fixation dot, the yellow dot is the target, and the four red dots represent the landmark. In the landmark shift condition, the pink dots represent a shifted landmark position.

The experiment consisted of 3 conditions: one with no-landmark (1), one with a stable landmark (2), and a cue-conflict condition with a landmark shift (3) (Figure 2C). A trial began when one of the three LEDs on the hand position bar illuminated, indicating the required initial hand position. The monkey then maintained hand fixation on that LED for 700 ms. The fixation dot appeared for 300 ms to allow the animal to fixate its gaze at a controlled location for each trial. In conditions two and three, the landmark and a fixation dot appeared for 500 ms. The target flashed for 100 ms, followed by a 300 ms delay displaying only the central fixation dot and landmark. A bright LED mask that faded to black near the edges covered the entire screen for 200 ms. The mask concealed the apparent motion of the landmark and prevented the monkey from shifting its attention to the landmark. It also prevented the influence of any aftereffects caused by the illumination of the target and landmark and ensured the monkey responded based on their memory rather than the afterimage of the target (Chen et al., 2014). The landmark and fixation dot appeared again for 200-600 ms, but in the cue-conflict condition, the landmark shifted by 8 ° in one of eight directions (up, down, left, right, and diagonally). Finally, the monkey had 2000 ms to respond by reaching and touching the screen where they thought the target was relative to the landmark, similar to Byrne and Crawford (2010). They received a reward when they touched within a 5.2 cm radius circle touch window. This window was large enough to include both the original target location and the newly shifted target location. The same was shown in the no-landmark condition except in the absence of a landmark. The initial gaze fixation in the no-landmark condition was also 500 ms longer. During this 500 ms, the landmark was displayed in the stable and landmark shift conditions.

### Data Analysis

The voltages from the head and eye coil were sampled at 1000 Hz, digitized, and converted into 2D angles of the eye and head in space. Experimental data were analyzed using MATLAB and the beginning and end of each saccade were marked manually.

To isolate the influence of the landmark shift from systematic reach errors, we subtracted the mean reach error (stable landmark condition) from the reach data in the landmark shift condition. This correction was performed for each of the 15 targets. In the no-landmark condition for M2, the three leftmost target locations were excluded from analysis because the animal consistently struggled to reach these targets. The remaining data were retained. Gaze onset times ≤ 0 ms (anticipatory saccades initiated at or before the go signal) were retained in all analyses and are reported as measured; the proportion of such trials was 11.2%, 22.2%, and 27.8% for the no-landmark, stable landmark, and landmark shift conditions, respectively, in M1, and 6.0%, 11.4%, and 8.6% in M2.

We used a combination of parametric and non-parametric tests, including Kruskal-Wallis tests with Dunn’s post-hoc comparisons, one-way ANOVAs, unpaired and paired t-tests, Wilcoxon signed-rank tests, and Fisher’s r-to-z tests, to test whether task condition and delay duration affected gaze onset time, gaze landing time, hand onset time, completion time, reach accuracy, reach variance, and gaze-hand temporal coupling. The influence of landmark shift direction on reach accuracy and variance was assessed using one-way ANOVAs and paired t-tests. The influence of target location and landmark distance on gaze and touch endpoints was assessed using Spearman correlations. Allocentric weights were compared across shift directions using Mann-Whitney U tests with Bonferroni correction, and their overall significance was tested using Wilcoxon signed-rank tests. No outliers were removed from the dataset. Statistical significance was set at p < 0.05, and Bonferroni correction was applied to post-hoc pairwise comparisons to control for multiple comparisons. All statistical tests were conducted using a combination of MATLAB, Prism 8, and Microsoft Excel.

## RESULTS

For the following analyses, we used 6228 trials (1774 no-landmark, 1513 stable landmark, 2941 landmark shift) from M1 and 6835 trials (2528 no-landmark, 1992 stable landmark, 2315 landmark shift) from M2. In the first two sections we examine the influence of landmarks and landmark shifts in the temporal and spatial domains, employing standard measures in the eye-hand coordination field. In the final section we examine ego-allocentric weighting and some of the influencing factors.

### Temporal Analyses

We first examined the time of four events (gaze onset, hand onset, gaze offset, hand touch) relative to the time when the fixation dot turned off (the ‘go’ signal). Figure 3A shows the mean (± 95% CI) horizontal gaze trajectory relative to the go signal (blue line), as well as the mean (± 95% CI) hand onset (black line) and offset (orange line) times for all trials toward one example target from each animal in the no-landmark condition. In these examples, saccades undershot the target (accuracy is quantified in a later section), and a single saccade step was used. In cases where multi-step saccades were used, both initial and final saccade end points were marked manually (Fig. 5). Hand motion was monitored online and appeared to follow continuous trajectories in both animals.

**Figure 3.**
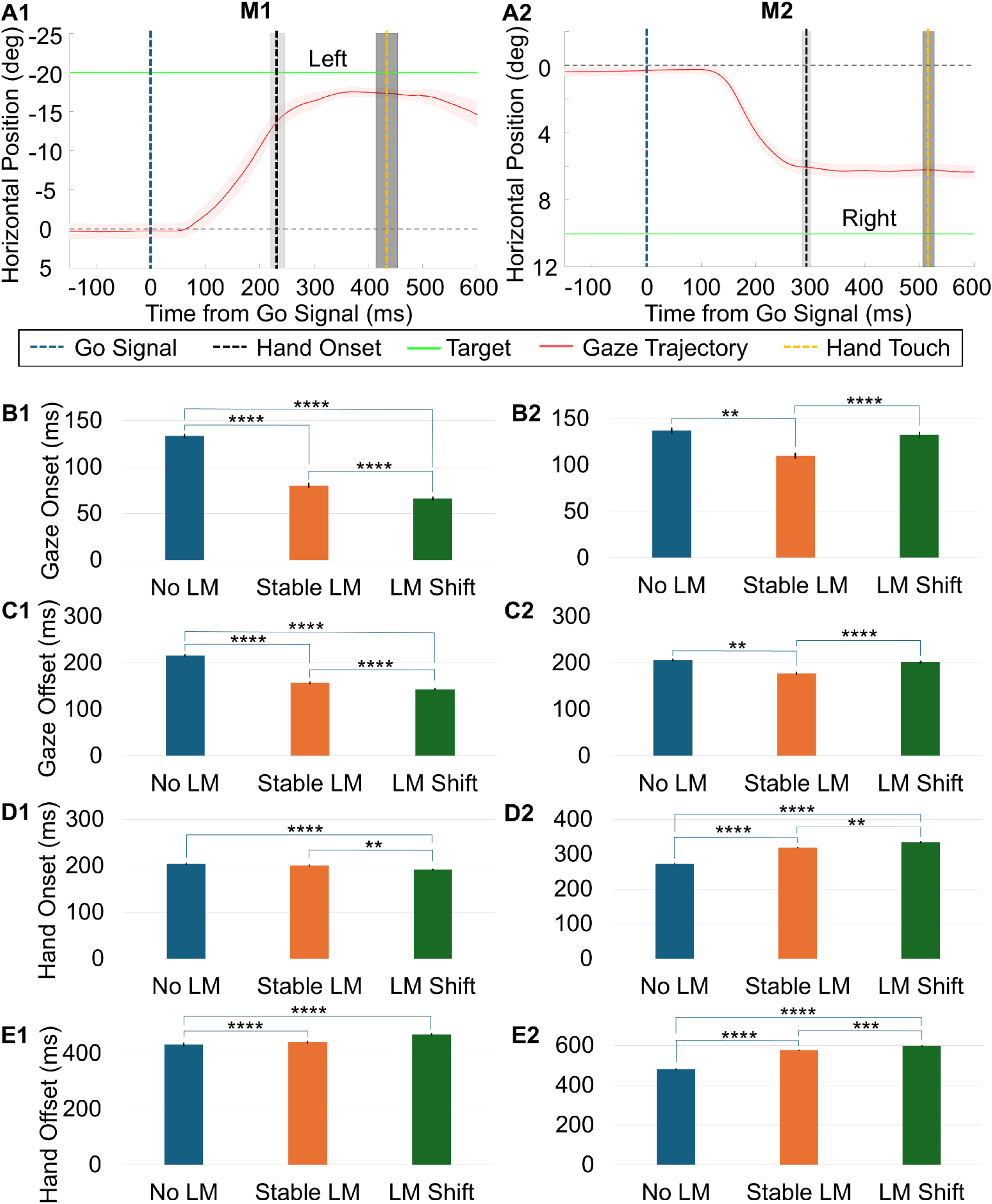
Example gaze trajectories and mean temporal measures for M1 (left column) and M2 (right column). (A) Averaged horizontal eye-in-space trajectory, hand onset times, and touch times for one example target ((−20, 0) for M1 and (10, 0) for M2) in the no-landmark condition, plotted as a function of time from the go signal (± 95% confidence intervals shown as shaded region). Time 0 ms represents home fixation extinction, signalling the animal to reach. (B) Mean gaze onset time by condition. (C) Mean gaze offset time by condition. (D) Mean hand onset time by condition. (E) Mean hand offset time by condition. Error bars show ± SEM. Trial counts per condition: M1: No LM: n = 1774, stable LM: n = 1513, LM shift: n = 2941; M2: no LM: n = 2528, stable LM: n = 1992, LM shift: n = 2315. Asterisks indicate significant pairwise differences (Dunn’s post-hoc test, Bonferroni-corrected). * p < 0.05, ** p < 0.01, *** p < 0.001, **** p < 0.0001.

Figure 3 B-E shows mean temporal latencies relative to the ‘go’ signal (across all trials and targets) in both animals, and Table 1 provides statistical comparisons across conditions (Kruskal-Wallis test followed by Dunn’s post-hoc test). Mean saccade durations were relatively consistent (M1: 78.76 ± 14.26 ms, M2: 68.78 ± 12.21 ms). Both gaze onset (Fig. 3B; time when gaze exceeded 50°/s) and offset times (Fig. 3C) were significantly higher in the no-landmark condition than in the landmark conditions for both animals, except for the landmark shift condition in M2 (gaze onset and gaze offset). The influence of landmarks on hand onset time (Fig 3D; time hand left LED bar) was inconsistent: lower in M1 but higher in M2. However, hand offset time (Fig. 3E; time of screen touch) was consistently later in the presence of landmarks. Overall, hand onset times were sometimes similar to gaze offset (M1, no-landmark condition) but in most cases occurred after gaze offset and of course lasted much longer (M1: 253.25 ± 253.10 ms, M2: 242.27 ± 83.56 ms).

**Table 1.**
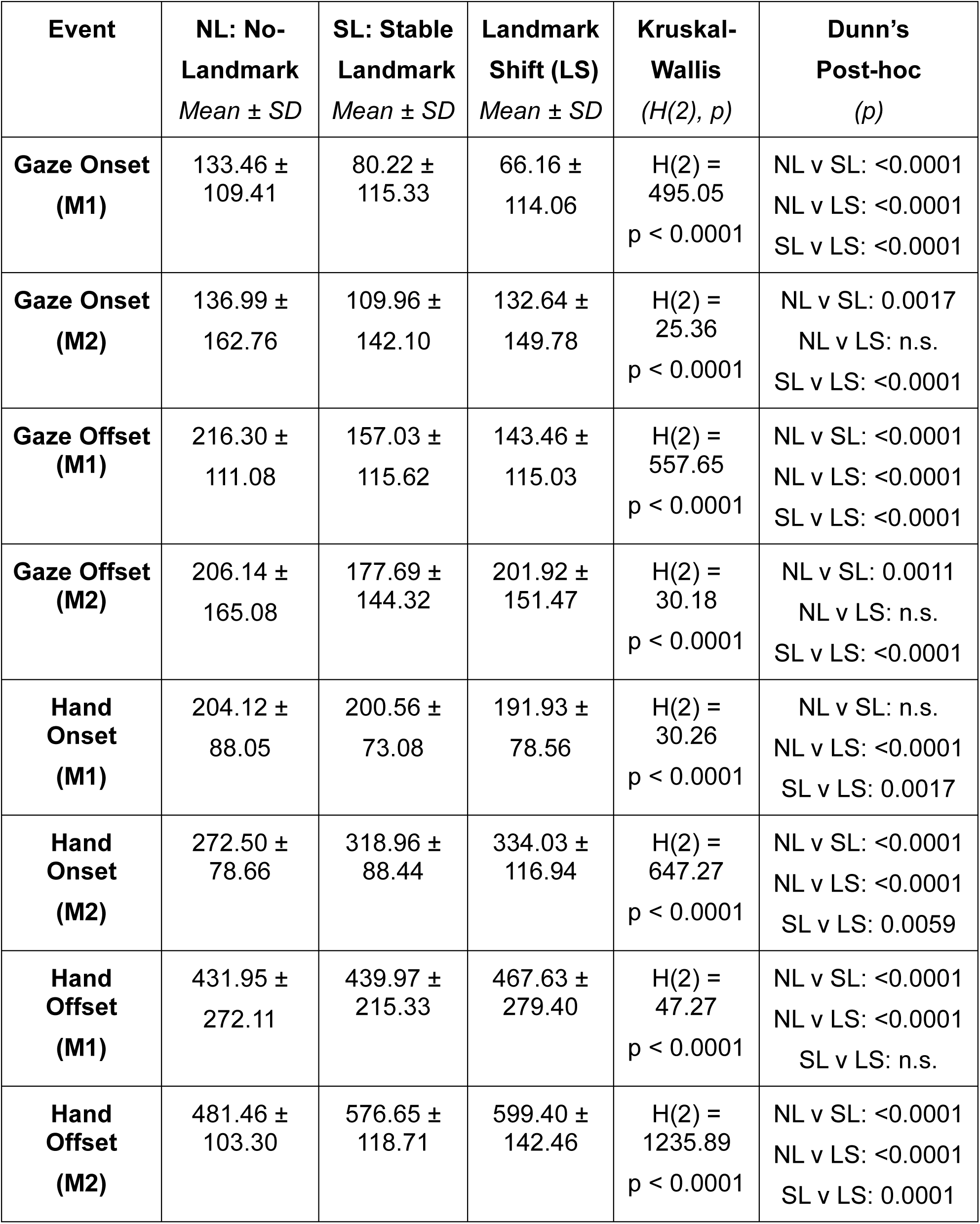
Mean temporal latencies (Mean ± SD, in ms) relative to the go signal.

INFLUENCE OF DELAY DURATION. To examine the effect of variable delay duration on gaze onset time, trials were divided into five ∼80 ms delay bins (198–278, 279–359, 360– 439, 440–520, and 521–600 ms), and both gaze and hand onset times were compared across bins within each condition for both animals (Figure 4).

**Figure 4.**
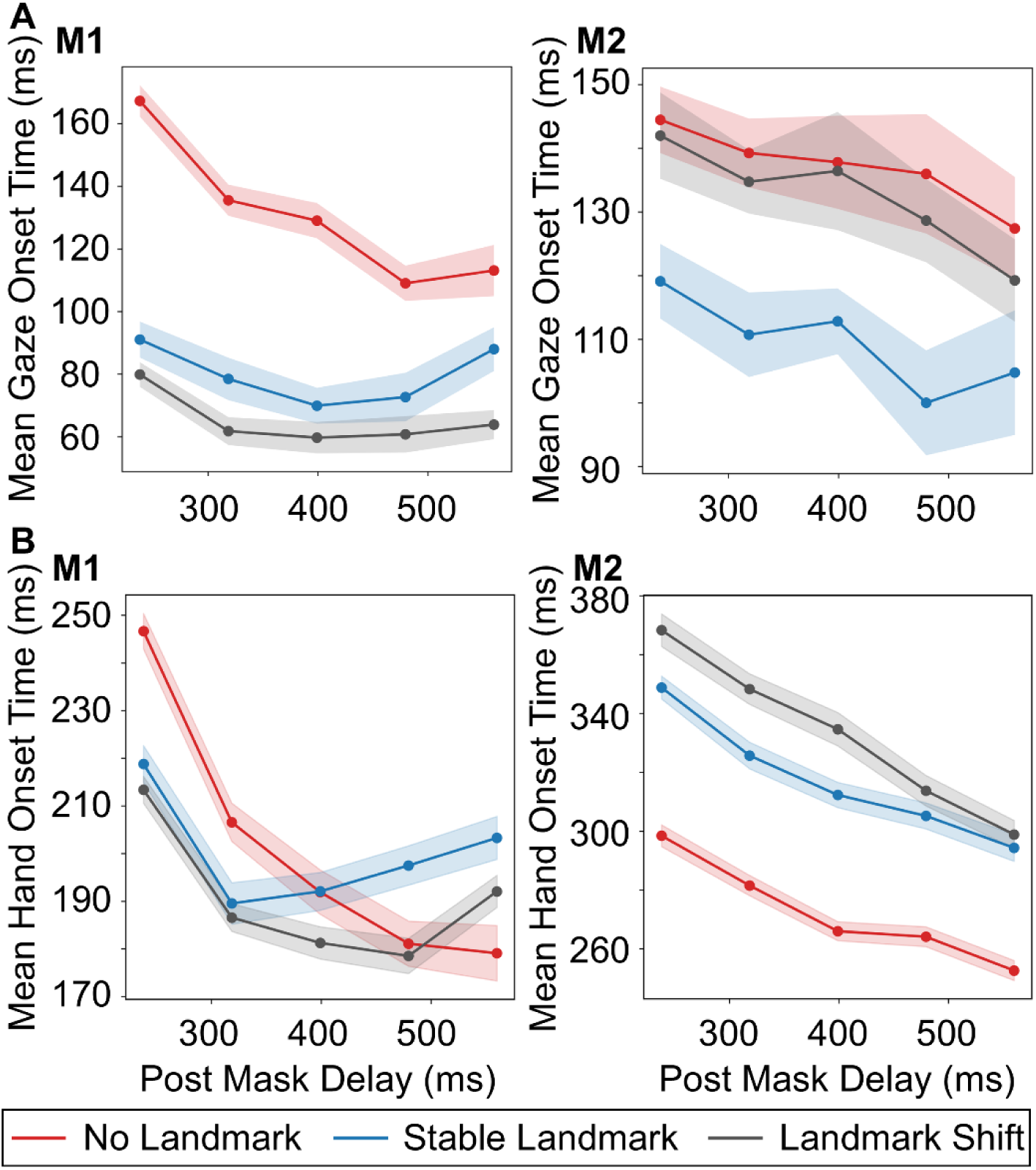
Mean gaze onset (A) and hand onset time (B) as a function of post-mask delay for M1 and M2. Trials were divided into five delay bins (198–278, 279–359, 360–439, 440–520, and 521–600 ms), and mean reaction time was computed within each bin for the no-landmark (red), stable landmark (blue), and landmark shift (grey) conditions. Points show the bin means, and shaded regions indicate ± SEM. Delay bin had a significant overall effect on gaze onset time in every condition except stable landmark in M2, and on hand onset time in every condition (Kruskal-Wallis).

In the case of gaze (Figure 4A), the variable delay period significantly affected onset time in most conditions (Kruskal-Wallis, all significant H(4) > 13, all p ≤ 0.0080), with the exception of the stable landmark condition in M2 (H(4) = 2.38, p = 0.6664). In all significant conditions, the shortest delay bin (198–278 ms) produced the longest gaze onset times and differed significantly from at least two longer bins in every condition except the M1 stable landmark condition, where it differed only from 360-439 ms bin (Dunn’s post-hoc, Bonferroni-corrected, all p ≤ 0.0417). For M1, the effect was most pronounced in the no-landmark condition, where gaze onset decreased consistently across bins (167.29 ± 101.42 ms to 113.11 ± 130.65 ms; Mean ± SD), with the shortest bin differing from all longer bins (all p < 0.0001) and the second bin also differing from the fourth (279–359 vs 440–520 ms, p = 0.0123). In the stable and landmark shift conditions of M1, significant effects were weaker and limited to comparisons involving the shortest or longest bins. For M2, significant effects were found in the no-landmark and landmark shift conditions, both driven primarily by differences between the shortest bin and longer bins, while the stable landmark condition showed no significant effect of delay on gaze onset.

For the hand (Figure 4B), the variable delay period had a significant effect on onset time across all task conditions in both animals (Kruskal-Wallis, all H(4) > 36, all p < 0.0001). In all conditions and both animals, the shortest delay bin (198–278 ms) produced the largest hand onset times and differed significantly from every longer delay bin (Dunn’s post-hoc, Bonferroni-corrected, all p ≤ 0.0292). Beyond this, the pattern varied. In M1, the most pronounced drop occurred between the shortest bin and the next (279–359 ms), after which hand onset times plateaued or showed no consistent trend, particularly in the stable landmark condition, where no significant differences were found among the three longest delay bins. In the no-landmark condition of M1, the decrease was more gradual and extended across bins (246.68 to 179.10 ms), with the second bin also differing significantly from the two longest bins (279–359 vs 440–520 ms, p = 0.0011; 279–359 vs 521–600 ms, p = 0.0007). In M2, hand onset time decreased more consistently with increasing delay across all conditions (no-landmark: 298.44 to 252.62 ms; landmark shift: 368.30 to 298.86 ms; stable: 348.82 to 294.30 ms), with significant pairwise differences observed across multiple bins beyond the shortest. Notably, in the landmark shift condition, M1 hand onset time was slightly elevated at the longest delay bin (521–600 ms) relative to the preceding bin (440–520 vs 521–600 ms, p = 0.0175), suggesting the relationship between delay and hand onset time is not strictly monotonic in this condition. Overall, reaction times tended to decrease with increasing delay for both gaze and hand onset, although this effect was more consistent for hand onset than for gaze onset.

GAZE-HAND TEMPORAL CORRELATIONS. To examine whether gaze and hand onset times were temporally coupled, and whether this coupling was affected by task condition, Spearman correlations between gaze onset and hand onset were calculated for each condition and animal. Gaze and hand onset times were significantly correlated across all conditions in both animals (all p < 0.0001). The strength of coupling differed significantly across conditions (Fisher’s r-to-z tests, Bonferroni-corrected). For M1, coupling was strongest in the no-landmark condition (ρ(1772) = 0.7579) and significantly weaker in both the stable landmark (ρ(1511) = 0.4215; z = 15.47, p < 0.0001) and landmark shift conditions (ρ(2939) = 0.4591; z = 16.46, p < 0.0001). Stable and landmark shift conditions did not differ from each other (z = −1.48, p = 0.4209). A consistent pattern was observed in M2: no-landmark coupling (ρ(2526) = 0.3814) was significantly stronger than both stable landmark (ρ(1990) = 0.1439; z = 8.57, p < 0.0001) and landmark shift (ρ(2313) = 0.1535; z = 8.58, p < 0.0001), with no significant difference between the two landmark conditions (z = −0.32, p = 1.0000). These results indicate that the temporal coupling between gaze and hand onset is significantly weaker when a landmark is present, regardless of whether that landmark is stable or shifting.

### Spatial Analysis

#### GENERAL OBSERVATIONS

Figure 5 shows example gaze trajectories (blue), vectors from initial to final hand position (orange), gaze position at the end of the first saccade (green), gaze position at hand landing (purple), and target positions (black), for each of our three experimental conditions and both animals (M1 above, M2 below). The data is provided in visual angles projected onto touch screen coordinates. Animals made centrifugal gaze shifts toward the targets in each of the three conditions, with a general tendency to overshoot in the upward direction. Hand trajectories started from three initial positions and converged toward the target array (only rewarded movements are shown). Smaller head movements toward the targets (not shown) were similar to those reported previously (Arora et al. 2019), and so are not repeated here. The following sub-sections describe the variable error, constant error, and gaze-hand correlations in these data.

**Figure 5.**
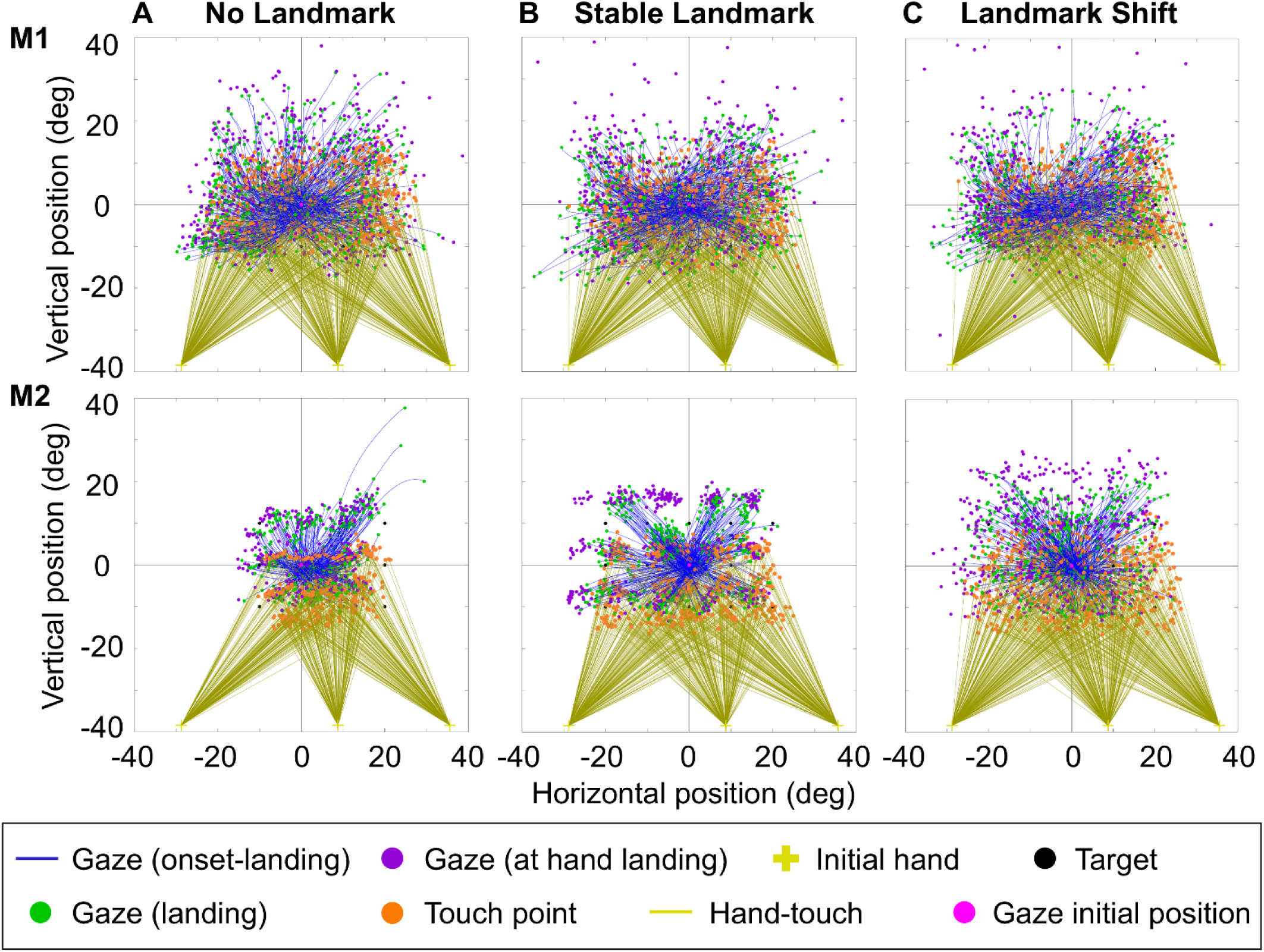
Example Gaze and touch endpoints, and 2-D gaze trajectories for the three visual conditions. Trajectories are plotted from the time of initial gaze motion to the end of the saccade. Complete gaze trajectories are provided, whereas the hand trajectories are only shown as vectors joining the initial and final hand positions. For M2, targets at the leftward positions were excluded in the no-landmark condition, as described in the Methods.

#### TWO-DIMENSIONAL ERROR ANALYSIS

To quantify the accuracy and precision of both gaze and reach movements, we fitted 95% confidence ellipses to the endpoint distributions around each target. For this analysis, we only used the targets that fell within the four-landmark array. Figure 6 shows the gaze (left column) and touch (2^nd^, 3^rd^ columns) endpoints (15 per target) for an example experiment, along with the 95% confidence ellipses fit to these data for M1 (see Supplemental Figure 1 for a similar plot for M2). The rows separate the three conditions by row (no-landmark, stable landmark, landmark shift), with space coordinates used in the first two columns and hand touch also shown in landmark coordinates (i.e., relative to the center of the four initial landmark cues) in the right column. (Note these were not present in the no-landmark condition, so we used the same spatial location for comparison). The individual endpoints for the initial gaze position are shown as green dots, and the gaze/hand end positions as blue dots. As expected, most of the data points are contained within the ellipses, and the touch data mostly falls within the landmark array (right column). In these example data, the gaze ellipses (left column) are much larger than the touch ellipses (middle column).

**Figure 6.**
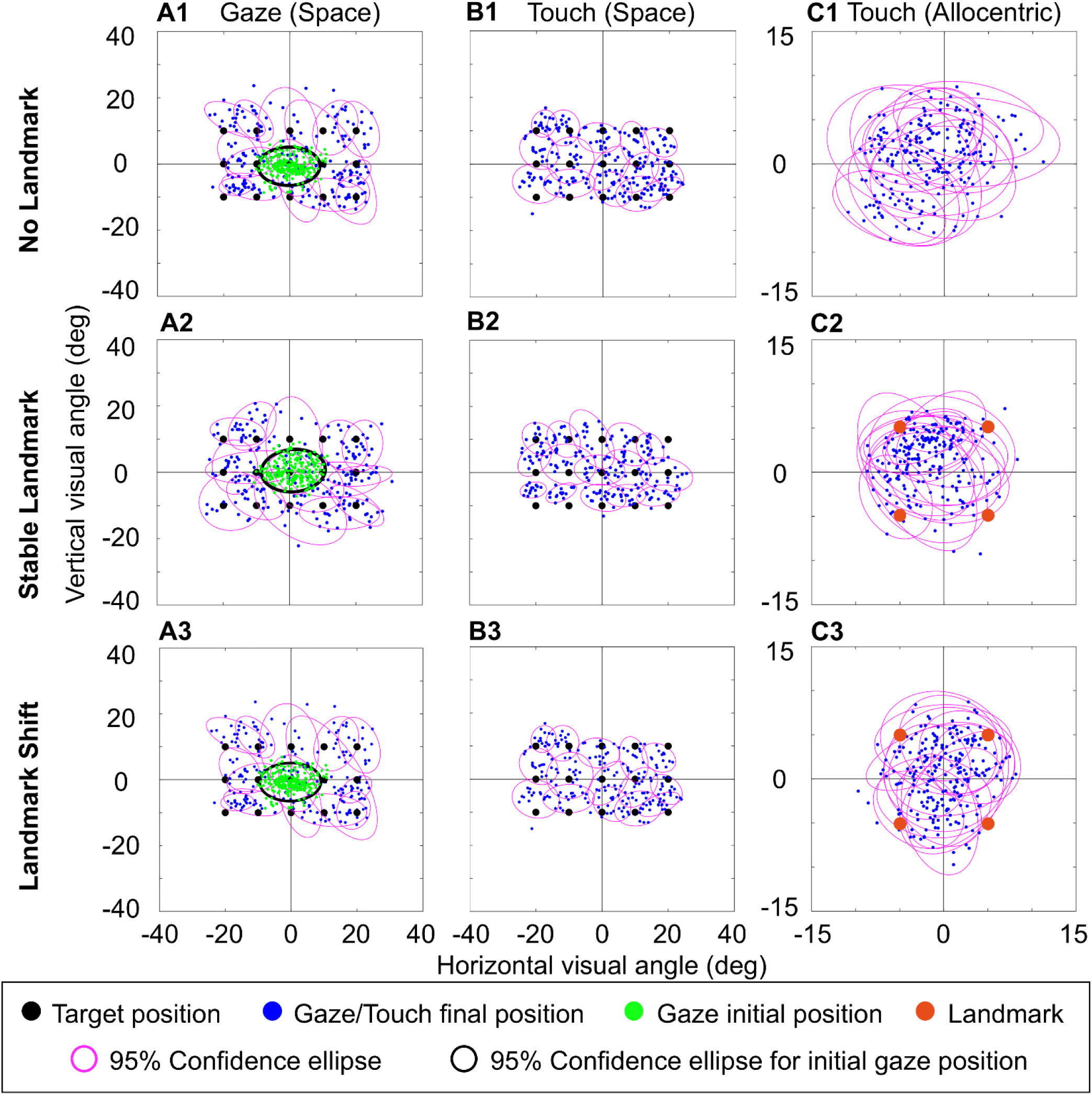
Example 95% confidence ellipses for data collected in one session in M1 (See Supplemental Figure 1 for M2). (A) gaze endpoints at the end of saccades in each condition (B) touch endpoints at the point of screen contact. (C) touch endpoints at the point of screen contact in allocentric coordinates.

Figure 7 A & B shows the 95% confidence ellipses fit to all of the gaze and touch endpoints collected from M1 and M2, plotted in space coordinates using the same conventions as Figure 6 (but without the original data points). Supplemental Figure 2 shows the corresponding plots in allocentric coordinates, which essentially shows that the data (excluding catch trials) consistently fell within the range of the four landmarks. The centers of these ellipses provided a measure of accuracy, whereas the areas provided a measure of precision. These measures are quantified in the following sub-sections.

**Figure 7.**
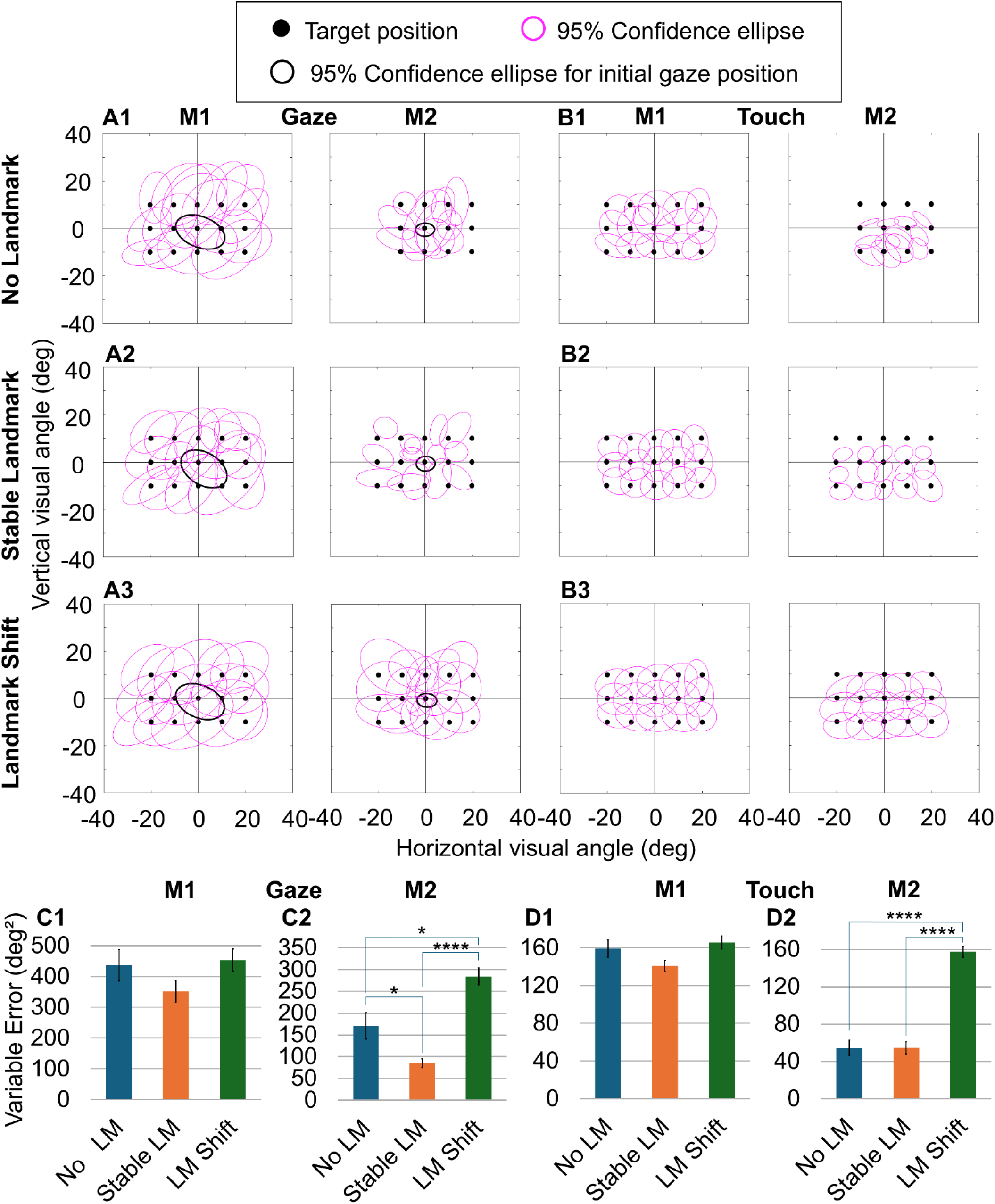
95% confidence ellipses fit to complete data sets of M1 and M2, plotted in visual coordinates. (A) Pink ellipses show the 95% confidence ellipses for gaze position at the end of saccades in each condition. (B) 95% confidence ellipses for touch endpoints at the point of screen contact. Black ellipses indicate the 95% confidence ellipse for initial gaze (fixation) position. The black dots indicate the target positions. (C) Bar graphs showing mean variable gaze error (ellipse area) for each condition for M1 and M2. (D) Bar graphs showing mean variable touch error (ellipse area) for each condition for M1 and M2. Error bars show ± SEM. Asterisks indicate significant pairwise differences (Dunn’s post-hoc test, Bonferroni-corrected). * p < 0.05, ** p < 0.01, *** p < 0.001, **** p < 0.0001.

#### VARIABLE ERROR (PRECISION)

Fig. 7 C/D plots mean ellipse areas (across targets) for gaze and touch end points for all conditions in both animals. The corresponding numbers and statistical comparisons (Kruskal-Wallis test followed by Dunn’s post-hoc test) between conditions are provided in Table 2. Across the three task conditions, gaze (in both animals) and touch (in M1) tended to be least variable in the stable landmark condition. In M2, gaze was significantly less variable in the stable landmark condition than in both the no-landmark and landmark shift conditions, whereas touch was significantly more variable in the landmark shift condition than in both the no-landmark and stable landmark conditions (Table 2). In M2, gaze was significantly more variable in the landmark shift condition than no-landmark. Overall, these data suggest fixed visual landmarks tend to stabilize aiming, whereas shifting landmarks destabilize aiming.

**Table 2.**
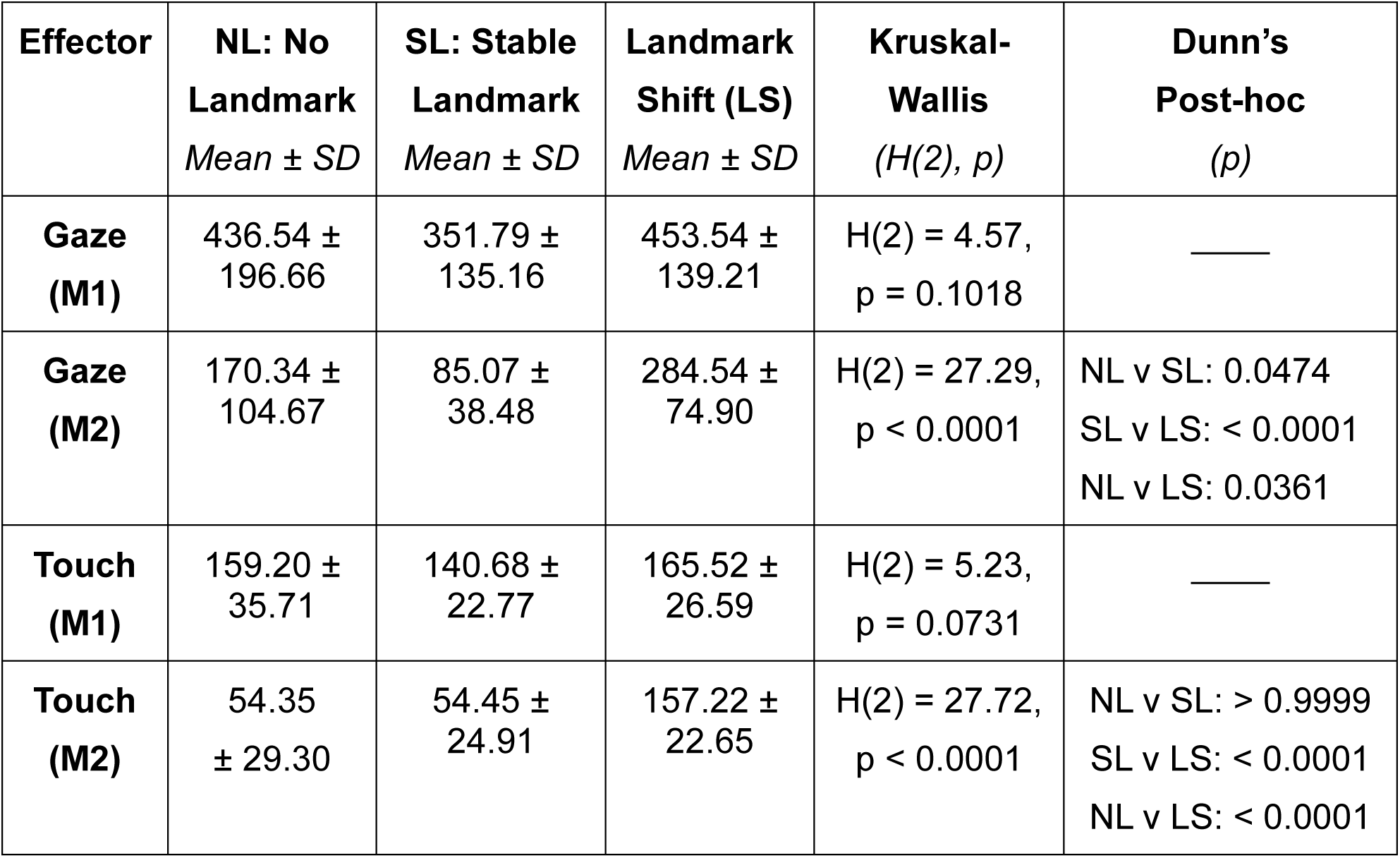
Variable error (degrees^2^ ellipse area) by task condition for M1 and M2.

| Effector | NL: No Landmark<br>*Mean ± SD* | SL: Stable Landmark<br>*Mean ± SD* | Landmark Shift (LS)<br>*Mean ± SD* | Kruskal-Wallis<br>*(H(2), p)* | Dunn’s Post-hoc<br>*(p)* |
|---|---|---|---|---|---|
| **Gaze (M1)** | 436.54 ± 196.66 | 351.79 ± 135.16 | 453.54 ± 139.21 | H(2) = 4.57, p = 0.1018 | ____ |
| **Gaze (M2)** | 170.34 ± 104.67 | 85.07 ± 38.48 | 284.54 ± 74.90 | H(2) = 27.29, p < 0.0001 | NL v SL: 0.0474<br>SL v LS: < 0.0001<br>NL v LS: 0.0361 |
| **Touch (M1)** | 159.20 ± 35.71 | 140.68 ± 22.77 | 165.52 ± 26.59 | H(2) = 5.23, p = 0.0731 | ____ |
| **Touch (M2)** | 54.35<br>± 29.30 | 54.45 ± 24.91 | 157.22 ± 22.65 | H(2) = 27.72, p < 0.0001 | NL v SL: > 0.9999<br>SL v LS: < 0.0001<br>NL v LS: < 0.0001 |

A general observation is that gaze ellipses (Fig. 7A, C) were much larger than touch ellipses (Fig. 7B, D). Gaze ellipse areas were significantly larger than the touch ellipses in all three conditions for M1 (two-tailed paired t-test: no-landmark t(14) = 6.086, p < 0.0001; stable landmark t(14) = 6.430, p < 0.0001; landmark shift t(14) = 8.411, p < 0.0001). For M2, the gaze ellipse areas were also significantly larger than the touch ellipses in all three conditions (two-tailed paired t-test: no-landmark t(11) = 4.045, p = 0.0019; stable landmark t(14) = 2.911, p = 0.0114; landmark shift t(14) = 6.058, p < 0.0001). This is consistent with the fact that we only rewarded touch accuracy, and that gaze tends to become less precise after a memory delay (Arora et al., 2019; Flanagan et al., 2008).

#### CONSTANT ERROR (ACCURACY)

This measure proved to be less consistent for task or effector. Significant task differences were only observed in M2 touch (H(2) = 7.16, p = 0.028), where constant error was largest for the no-landmark condition and significantly lower in the landmark shift condition (p = 0.023). Further, a significant difference between gaze and hand touch was only observed in the landmark shift condition for M1 (two-tailed paired t-test, t(14) = 2.638, p = 0.0195). Similar results were observed when error was separated into horizontal and vertical components.

#### INFLUENCE OF MEMORY DELAY ON REACH ACCURACY AND PRECISION

To examine whether the memory delay duration influenced reach errors (as it did on reaction time), we separated data from the same five delay bins used in our temporal analysis. The variable delay period had no significant effect on mean Euclidean error in five of the six animal-by-condition cases (Kruskal-Wallis, all H(4) ≤ 7.63, all p > 0.10; Supplemental Figure 5). The exception was M2 in the stable landmark condition (H(4) = 18.37, p = 0.0010), where error was lowest in the longest delay bin (521–600 ms). Delay also had no effect on constant error and variable error in either animal (all Kruskal-Wallis H(4) < 3.50, all p > 0.48) (Supplemental Figure 6).

#### INFLUENCE OF TARGET POSITION

Figure 8 A, B show the correlation of gaze in space (left column), hand touch in space (middle column), and hand in landmark coordinates (right column) against horizontal target position in our three conditions (See Supplemental Figure 3 for a very similar plot for M2). Figure 8 D1,2 and Table 3 show the corresponding Spearman correlations for both animals. In every case, gaze and touch were significantly correlated with target position in space. Similarly significant, though slightly weaker, correlations were found for the more limited range of the 3 vertical target positions (Table 3).

**Figure 8.**
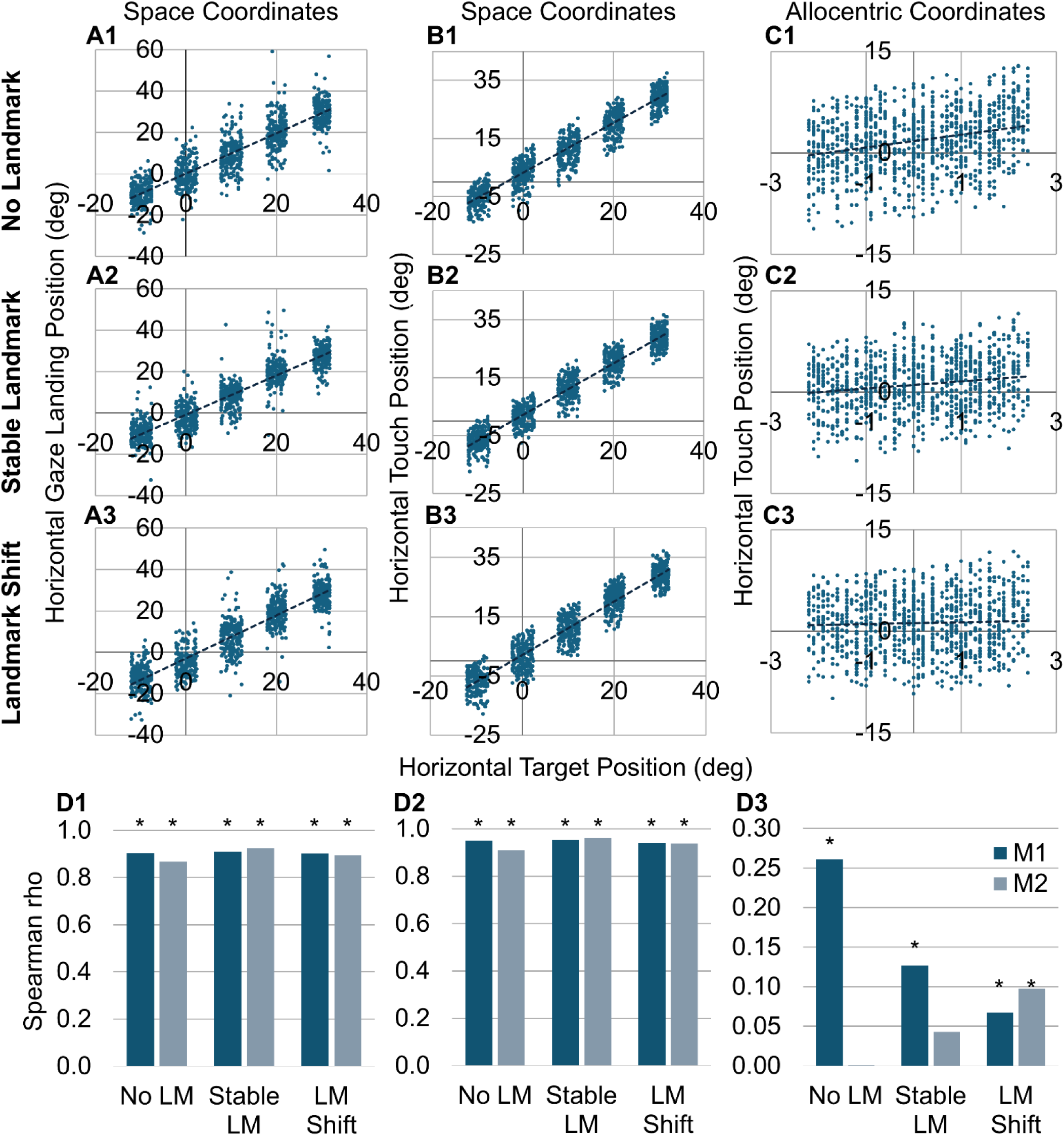
Correlation of gaze and hand position to target position in M1 (see Supplemental Figure 3 for M2), shown in the horizontal dimension. Each row represents a condition. (A) Correlation of gaze landing position to target position in space coordinates. (B) Correlation of hand touch position to target position in space coordinates. (C) Correlation of hand touch position to target position in allocentric (landmark-centered) coordinates. (D) Bar graphs summarizing the Spearman correlation coefficient for each condition for gaze (D1), touch (D2) and allocentric touch (D3), with M1 (dark blue) and M2 (light blue) shown side by side. Asterisks indicate significant correlations (p < 0.05).

**Table 3.**
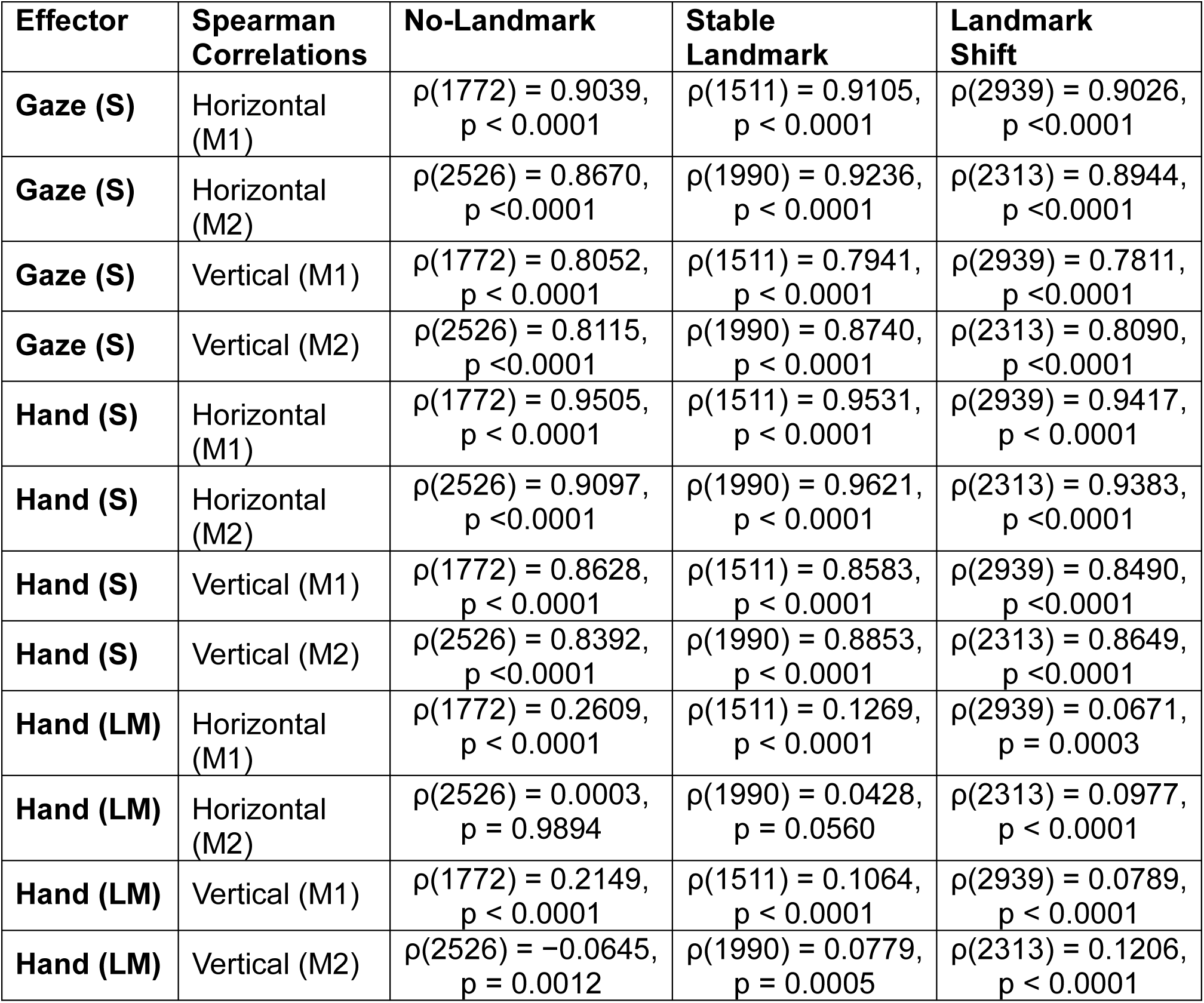
Spearman correlation coefficients (ρ) between gaze and touch position versus target position in space (S) or Landmark (LM) -centered coordinates.

| **Effector** | **Spearman Correlations** | **No-Landmark** | **Stable Landmark** | **Landmark Shift** |
|---|---|---|---|---|
| **Gaze (S)** | Horizontal (M1) | ρ(1772) = 0.9039, p < 0.0001 | ρ(1511) = 0.9105, p < 0.0001 | ρ(2939) = 0.9026, p <0.0001 |
| **Gaze (S)** | Horizontal (M2) | ρ(2526) = 0.8670, p <0.0001 | ρ(1990) = 0.9236, p < 0.0001 | ρ(2313) = 0.8944, p <0.0001 |
| **Gaze (S)** | Vertical (M1) | ρ(1772) = 0.8052, p < 0.0001 | ρ(1511) = 0.7941, p < 0.0001 | ρ(2939) = 0.7811, p <0.0001 |
| **Gaze (S)** | Vertical (M2) | ρ(2526) = 0.8115, p <0.0001 | ρ(1990) = 0.8740, p < 0.0001 | ρ(2313) = 0.8090, p <0.0001 |
| **Hand (S)** | Horizontal (M1) | ρ(1772) = 0.9505, p < 0.0001 | ρ(1511) = 0.9531, p < 0.0001 | ρ(2939) = 0.9417, p < 0.0001 |
| **Hand (S)** | Horizontal (M2) | ρ(2526) = 0.9097, p <0.0001 | ρ(1990) = 0.9621, p < 0.0001 | ρ(2313) = 0.9383, p <0.0001 |
| **Hand (S)** | Vertical (M1) | ρ(1772) = 0.8628, p < 0.0001 | ρ(1511) = 0.8583, p < 0.0001 | ρ(2939) = 0.8490, p <0.0001 |
| **Hand (S)** | Vertical (M2) | ρ(2526) = 0.8392, p <0.0001 | ρ(1990) = 0.8853, p < 0.0001 | ρ(2313) = 0.8649, p <0.0001 |
| **Hand (LM)** | Horizontal (M1) | ρ(1772) = 0.2609, p < 0.0001 | ρ(1511) = 0.1269, p < 0.0001 | ρ(2939) = 0.0671, p = 0.0003 |
| **Hand (LM)** | Horizontal (M2) | ρ(2526) = 0.0003, p = 0.9894 | ρ(1990) = 0.0428, p = 0.0560 | ρ(2313) = 0.0977, p < 0.0001 |
| **Hand (LM)** | Vertical (M1) | ρ(1772) = 0.2149, p < 0.0001 | ρ(1511) = 0.1064, p < 0.0001 | ρ(2939) = 0.0789, p < 0.0001 |
| **Hand (LM)** | Vertical (M2) | ρ(2526) = −0.0645, p = 0.0012 | ρ(1990) = 0.0779, p = 0.0005 | ρ(2313) = 0.1206, p < 0.0001 |

It is possible that in the landmark conditions, correlations in space could arise from a strategy of pointing toward the centre of the landmark arrays. To ensure that this was not the case, we checked if there was still a correlation between hand position and target positions in a coordinate system based on the landmarks that surround the targets, i.e., where the centre of the square is zero (Figure 8C, D3, Table 3). (For comparison, we treated no-landmark trials as if the landmark was visible and used initial landmark positions for the shift condition.) In this coordinate system, horizontal target-touch correlations were modest, but mostly significant (the exception being M2 in the no-landmark and stable landmark conditions). Vertical target-touch correlations were modest and consistently significant (Table 3). One reason for these low correlations might be because of the very small target range relative to measurement noise in this coordinate system. To test this, we included catch trials where the target was located outside the landmark array. For these datasets, the correlation between touch and target position in landmark coordinates was substantially higher in both the horizontal (M1: Spearman’s ρ(1186) = 0.6134, p < 0.0001; M2: Spearman’s ρ(2057) = 0.5769, p < 0.0001) and vertical dimensions (M1: Spearman’s ρ(1186) = 0.6264, p < 0.0001; M2: Spearman’s ρ(2057) = 0.5855, p < 0.0001) for both animals. Overall, these data show that reaches did not simply aim toward the centre of the target array but were also modulated by target position.

#### GAZE-HAND COORDINATION

As noted in the introduction, humans and monkeys often show a tight correlation between final gaze and hand position (Johansson et al., 2001; Neggers & Bekkering, 2000). We found the same in a similar head-unrestrained task without a memory delay (Arora et al., 2019). The preceding analysis above suggests that this coupling might break down after a memory delay, due to reduced gaze accuracy. To examine this directly, we correlated horizontal/vertical gaze position as a function of hand position (Figure 9, Supplemental Figure 4, Table 4). Figure 9A1-3 plots final horizontal hand position as a function of final horizontal gaze position in space coordinates, for all three conditions. Figure 9 D1 shows the corresponding Spearman correlations, and Table 4 provides the correlations for both horizontal and vertical positions. All correlations were highly significant, suggesting some degree of eye-hand coupling (Arora et al. 2019).

**Figure 9.**
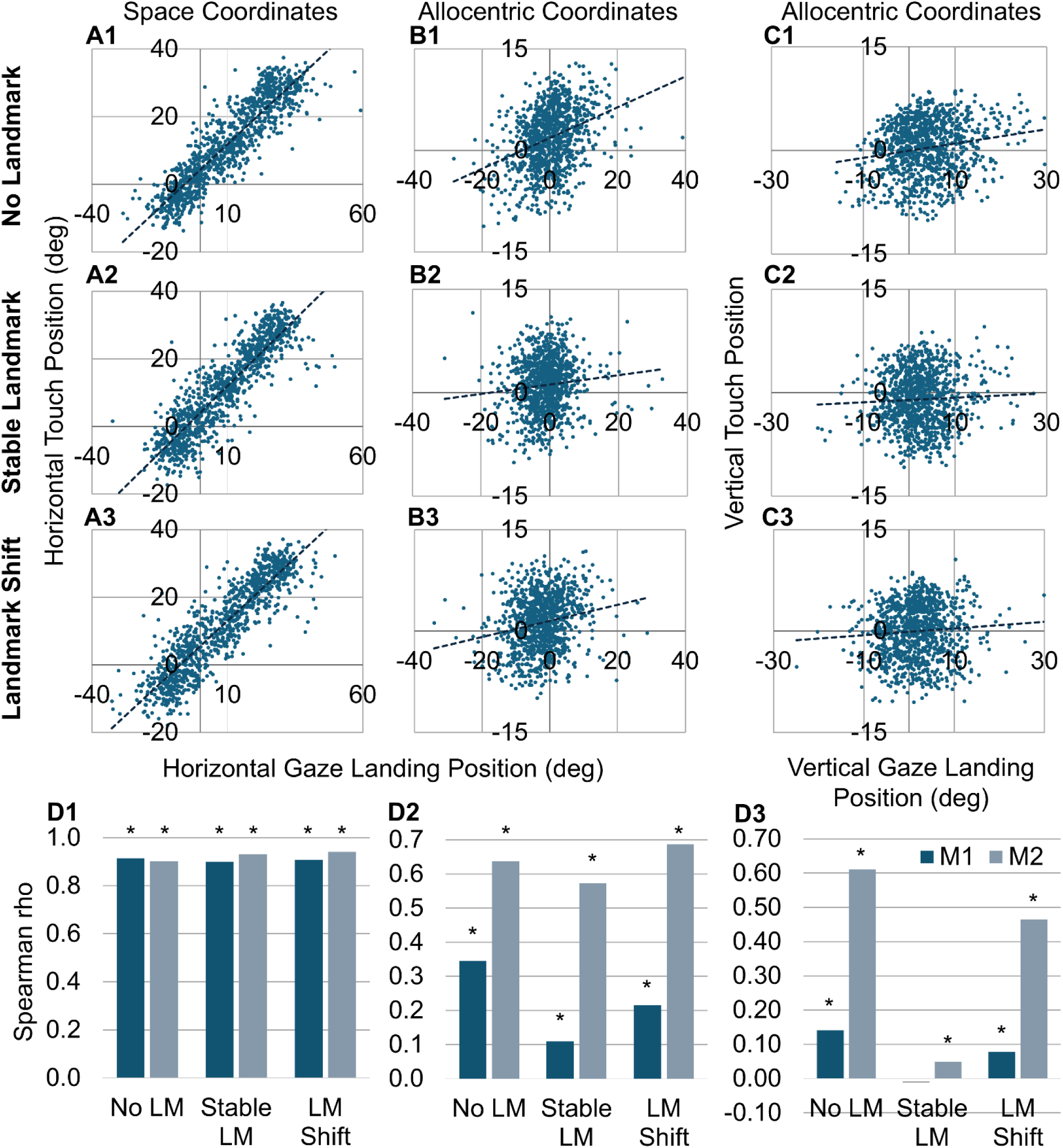
Correlation of hand touch position to gaze landing position in M1 (see Supplemental Figure 4 for M2). (A) Correlation of horizontal hand touch position to gaze position in space coordinates. (B) Correlation of horizontal hand touch to gaze position in allocentric (landmark-centered) coordinates. (C) Correlation of vertical hand touch to gaze position in allocentric coordinates. (D) Bar graphs summarizing the Spearman correlation coefficient of each condition for horizontal position in space (D1), and horizontal (D2) and vertical (D3) components in allocentric coordinates, with M1 (dark blue) and M2 (light blue) shown side by side. Asterisks indicate significant correlations (p < 0.05).

**Table 4.** Spearman correlation coefficients (ρ) between gaze versus touch position in space and landmark-centered coordinates.

| **Reference Frame** | **Spearman Correlations** | **No-Landmark** | **Stable Landmark** | **Landmark Shift** |
|---|---|---|---|---|
| **Space** | M1, Horizontal | ρ(1772) = 0.9146,<br>p < 0.0001 | ρ(1511) = 0.8997,<br>p < 0.0001 | ρ(2939) = 0.9077,<br>p < 0.0001 |
| **Space** | M2, Horizontal | ρ(2526) = 0.9030,<br>p < 0.0001 | ρ(1990) = 0.9314,<br>p < 0.0001 | ρ(2313) = 0.9404,<br>p < 0.0001 |
| **Space** | M1, Vertical | ρ(1772) = 0.7822,<br>p < 0.0001 | ρ(1511) = 0.7450,<br>p < 0.0001 | ρ(2939) = 0.7422,<br>p < 0.0001 |
| **Space** | M2, Vertical | ρ(2526) = 0.8192,<br>p < 0.0001 | ρ(1990) = 0.8580,<br>p < 0.0001 | ρ(2313) = 0.8789,<br>p < 0.0001 |
| **Landmark** | M1, Horizontal | ρ(1772) = 0.3452,<br>p < 0.0001 | ρ(1511) = 0.1096,<br>p < 0.0001 | ρ(2939) = 0.2147,<br>p < 0.0001 |
| **Landmark** | M2, Horizontal | ρ(2526) = 0.6367,<br>p < 0.0001 | ρ(1990) = 0.5727,<br>p < 0.0001 | ρ(2313) = 0.6871,<br>p < 0.0001 |
| **Landmark** | M1, Vertical | ρ(1772) = 0.1406,<br>p < 0.0001 | ρ(1511) = −0.0093,<br>p = 0.7164 | ρ(2939) = 0.0787,<br>p < 0.0001 |
| **Landmark** | M2, Vertical | ρ(2526) = 0.6109,<br>p < 0.0001 | ρ(1990) = 0.0487,<br>p = 0.0297 | ρ(2313) = 0.4649,<br>p < 0.0001 |

Presumably, many of these correlations that we observed in space coordinates could be due to the large variance in target / landmark positions in this coordinate system. When gaze and hand positions were expressed in landmark-centered coordinates (Figure 9B, 9C, 9D2, 9D3; Table 4), correlations were still significant in most cases, but markedly reduced. This was especially true in animal M1, and in the stable landmark condition (Figure 9 D2, D3). To test whether the difference in gaze-hand correlations across conditions were significant, we used Fisher r-to-z tests. In landmark-centered coordinates, gaze-hand correlations differed significantly between nearly all condition pairs in both animals (Bonferroni-corrected p < 0.05), the only exception was M1 vertical, no-landmark vs landmark shift. The most notable pattern was the reduction of the landmark-centered correlation in the stable landmark condition relative to no-landmark for both M1 (Horizontal: z = 7.136, p < 0.0001; Vertical: z= 4.306, p < 0.0001) and M2 (Horizontal: z = 3.371, p = 0.0008; Vertical: z= 22.069, p < 0.0001). This suggests that 1) eye-hand coupling is still present but degraded during memory-guided reaches, compared to immediate reaches (Arora et al. 2019), and 2) animals may rely less on spatial eye-hand coupling in the presence of a stable landmark.

### Landmark shift Direction and Allocentric Weight

#### INFLUENCE OF SHIFT DIRECTION

We saw that the landmark shift condition reduced gaze and reach accuracy relative to the original target positions when data are pooled across shift directions. Here, we examined if this influence is spatially specific to the direction of the shift, i.e., did gaze / reach endpoints shift in the same direction as the landmark? Past experiments suggest human reach endpoints and monkey gaze endpoints shift (partially) in the direction of the landmark shift (Byrne & Crawford, 2010; Li et al., 2017). This can be observed in Figure 10, which shows ellipses fit to the spatial distribution of reach endpoints to the same targets, but separated for leftward and rightward landmark shifts in both animals (Supplemental Figures 7 and 8 show similar plots for all shift directions). For M1, reach endpoints were displaced 2.49 ± 4.79° in the leftward direction on leftward-shift trials and 2.05 ± 4.62° in the rightward direction on rightward-shift trials (Mean ± SD). For M2, the displacements were larger: 3.60 ± 3.16° leftward and 4.20 ± 3.46° rightward. These endpoints were significantly displaced in the direction of the landmark shift for both shift directions in both animals (Wilcoxon signed-rank test, all p < 0.0001). (A means to normalize and comprehensively test these shifts is presented in the next section.)

**Figure 10.**
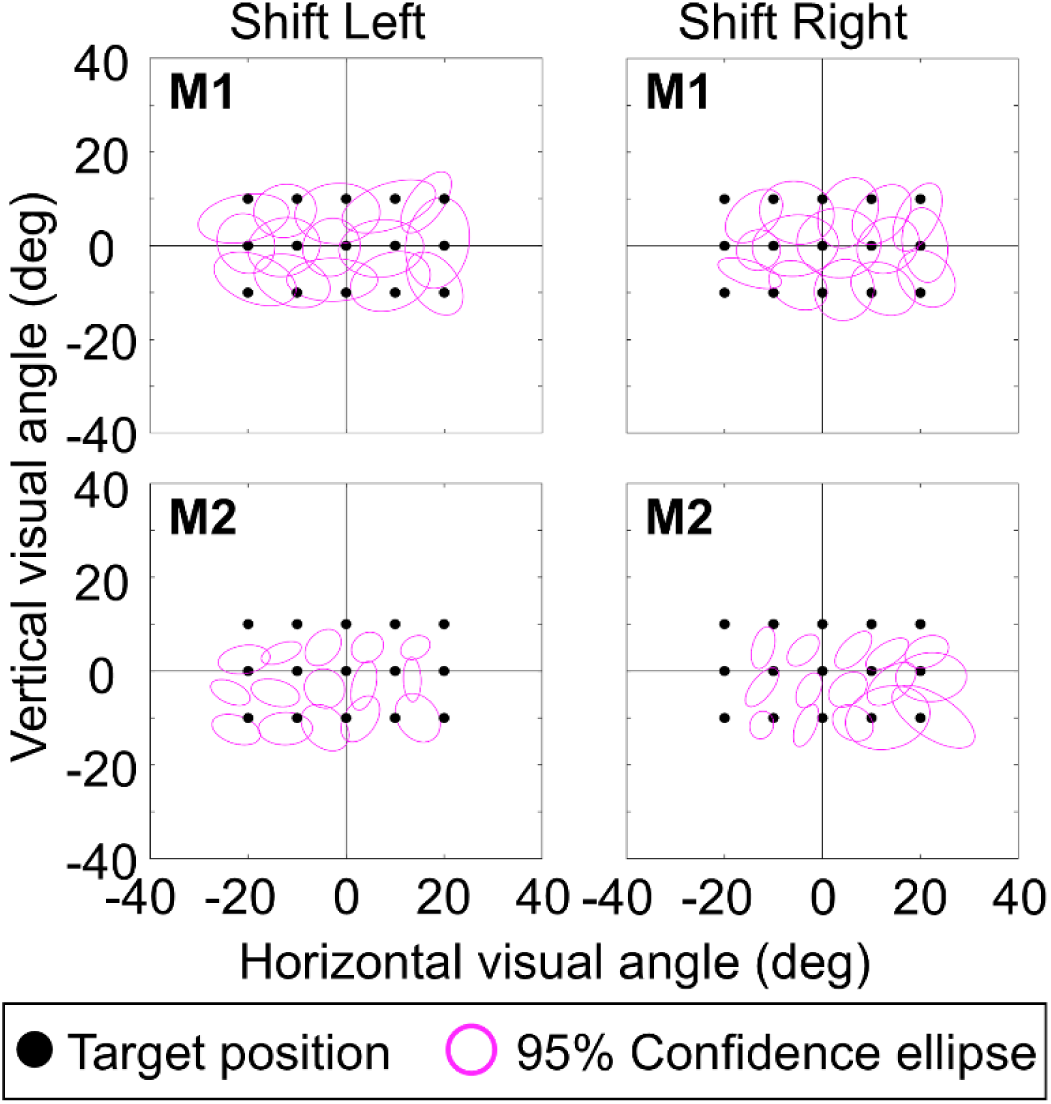
Reach endpoint distributions in the landmark shift condition for M1 (upper panels) and M2 (lower panels), separated by shift direction. Each panel shows 95% confidence ellipses for touch endpoints in a representative session, plotted in visual angle coordinates. Left column: trials in which the landmark shifted to the left; right column: trials in which the landmark shifted to the right. Black dots indicate target positions. The systematic displacement of ellipse centers in the direction of the landmark shift illustrates the allocentric influence on reach endpoints

Shift direction had a significant effect on touch accuracy for M1 (one-way ANOVA, F(7,112) = 3.77, p = 0.0010) but not for M2 (F(7,112) = 1.17, p = 0.3275). For M1, a landmark shift toward the upper left produced the highest accuracy (mean distance = 2.60 ± 0.94°; Mean ± SD) and was significantly more accurate than a shift toward the upper right (two-tailed paired t-test, t(14) = 3.27, Bonferroni-corrected p = 0.039), the right (t(14) = 3.71, corrected p = 0.016). Differences from the lower right (t(14) = 2.73, p = 0.11), and downward shifts (t(14) = 2.44, p = 0.20) do not remain significant after correction. Accuracy for the upper-left shift did not differ significantly from shifts toward the left, up, or lower left (all corrected p > 0.05). For M2, the most accurate direction was upward (mean distance = 3.38 ± 1.38°), though no shift direction differed significantly from any other. In contrast, shift direction did not significantly affect the area of the ellipses in either animal (M1: F(7,112) = 1.36, p = 0.2289; M2: F(7,112) = 1.40, p = 0.2118), indicating that shift direction did not affect the variance of touch responses in either monkey.

#### ALLOCENTRIC WEIGHTS

To quantify the influence of the landmark shift across all directions we calculated the allocentric weight (AW) parameter used in previous studies (Byrne & Crawford, 2010; Li et al., 2017). To do this, we first subtracted the mean touch/gaze error for each target in the stable landmark condition from each touch/gaze response for the same target in the landmark shift condition (to remove systematic egocentric biases in the monkey’s reach/gaze behaviour). The remaining error for each trial was then rotated into a coordinate system where the shift direction was to the right (Figure 11A), allowing one to ‘normalize’ and combine the effect across our entire dataset.

**Figure 11.**
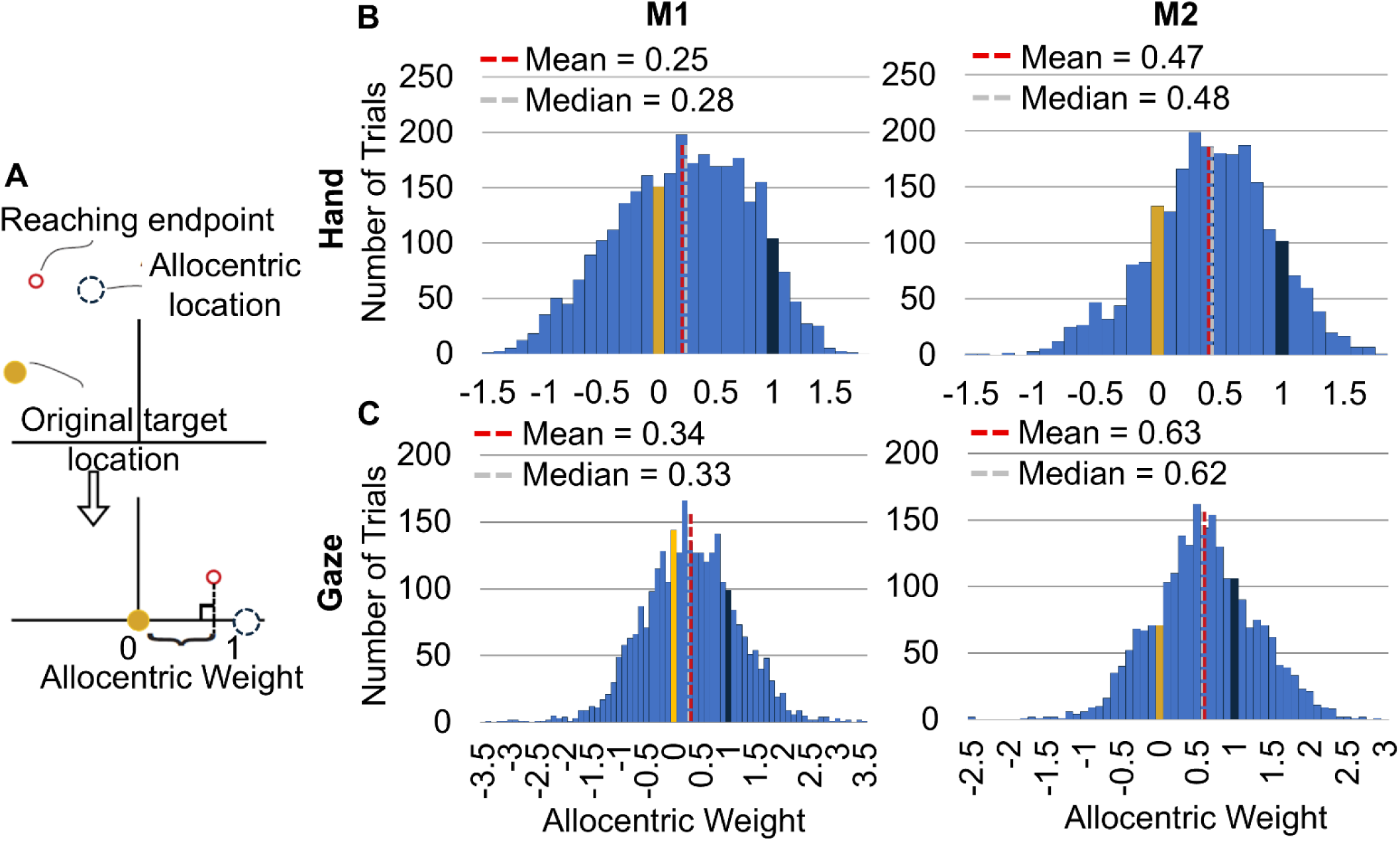
Distribution of allocentric weights in both animals (M1 and M2). (A) Coordinate system where the shift direction was to the right. (B) Histogram showing the distribution of allocentric weights for touch plotted against the number of trials (Y axis) collapsed across all sessions. (C) Histogram showing the distribution of allocentric weights for gaze plotted against the number of trials (Y axis) collapsed across all sessions.

Finally, the component of gaze / hand shift to the right of the target is divided by the amplitude of the landmark shift for each trial to compute the landmark influence.

Figure 11C shows the distribution of gaze endpoint errors in this rotated coordinate system. Although AW is theoretically bounded between 0 (purely egocentric) and 1 (purely allocentric), individual trial values can fall outside this range due to other sources of variable error in gaze, memory, and reach execution. For this reason means and medians of the resulting distributions are used to represent AW (Byrne & Crawford, 2010; Li et al., 2017). The gaze AW (M1: mean = 0.34, median = 0.33 / M2: mean = 0.63, median = 0.62) was significantly shifted toward the allocentrically defined target location T’ (M1: Wilcoxon Signed Rank Test, p < 0.0001/ M2: Wilcoxon Signed Rank Test, p < 0.0001).

Figure 11B shows similar distributions of the touch AW values (M1: mean = 0.25, median = 0.28/ M2: mean = 0.47, median = 0.48), which were also significantly shifted toward T’ (M1: Wilcoxon Signed Rank Test, p < 0.0001/ M2: Wilcoxon Signed Rank Test, p < 0.0001), indicating the influence of the shifted landmark on reaching endpoints. These weights were also consistent across different directions (e.g., in Figure 10: M1: left shifts 0.30, right shifts 0.26; M2: left shifts 0.51, right shifts 0.52). In short, although animal M2 showed a stronger AW for both gaze and reach than M1, the data for both animals fell within the intermediate ego-allocentric range expected from previous studies.

#### INFLUENCE OF GAZE ON REACH ‘AW’

A previous study found that allocentric weight of gaze shifts interacts with initial gaze direction (Li et al., 2017). To test this, first we examined the influence of landmark – gaze distance (Figure 12A) using landmarks in the same row or the same column as the initial gaze position. In both animals there was a trend for AW to drop for more peripheral landmarks, but the details and statistics were inconsistent. There was no significant effect of landmark distance on AW for M1 in either the horizontal (One-way ANOVA, F(2,997) = 0.33, p = 0.7199) or vertical dimension (two-tailed unpaired t-test, t(563) = 1.82, p = 0.0693). For M2, landmark distance had a significant effect on horizontal AW (One-way ANOVA, F(2,807) = 5.49, p = 0.0043; Bonferroni-corrected pairwise comparison 10° vs 20°, p = 0.003), but the pattern was not monotonic: AW was intermediate at 0° (0.5135), highest at 10° (0.5543), and then lowest at 20° (0.4367). There was no significant effect of landmark distance on vertical AW for M2 (two-tailed unpaired t-test, t(434) = 0.00, p = 0.9980).

**Figure 12.**
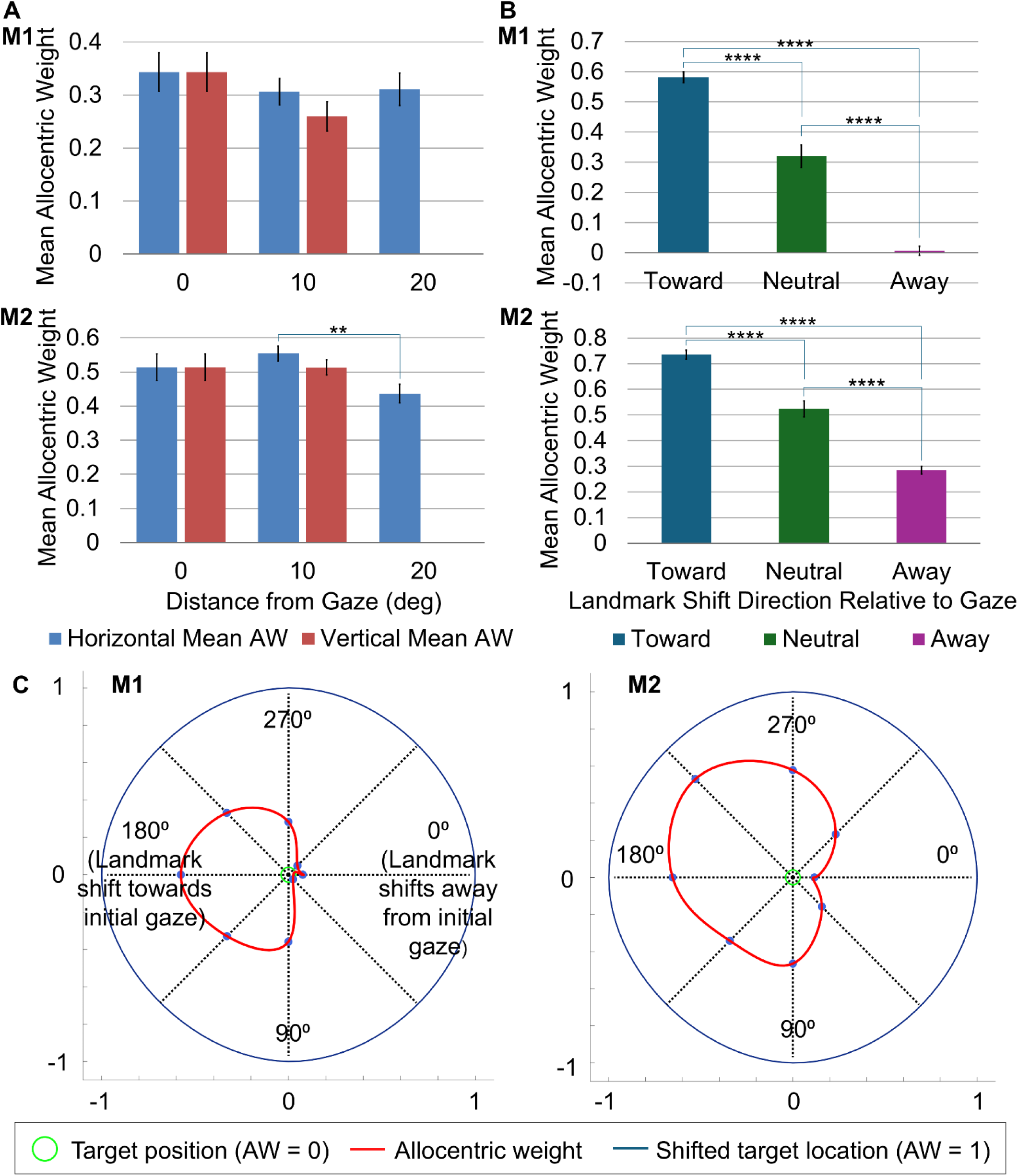
Influence of landmark shift direction and initial landmark distance from initial gaze position on allocentric weight (AW) in both animals (M1 and M2). (A) Comparison of the mean AW (Y axis) between landmarks divided into their horizontal and vertical positions located closer or further from the initial gaze position (X-axis). (B) Comparison of the mean AW (y-axis) between landmarks that shift toward, away from, or perpendicular to (neutral) the initial gaze position (X-axis). (C) Overall allocentric weight sorted by landmark shift direction relative to initial gaze. Error bars show ± SEM. Asterisks indicate significant pairwise differences (A: Bonferroni-corrected pairwise comparisons after one-way ANOVA; B: Dunn’s post-hoc test, Bonferroni-corrected). ** p < 0.01; **** p < 0.0001.

Second, we confirmed that reach AW is greater when landmarks shift toward, rather than away from gaze position (Li et al., 2017). This is shown in Figure 12B for data sorted into landmark shifts toward, neutral, or away from gaze, and in Figure 12C for all shift directions relative to gaze. For M1, the AW when the landmark shifted away from the initial gaze position was 0.007 and increased when the landmark shifted toward the initial gaze position (AW = 0.581). The AW was intermediate when the landmark shift was neutral (AW = 0.32). For M2, the away AW was 0.285, toward AW was 0.735, and neutral AW was 0.5238. The difference in AW due to landmark shift direction was highly significant for both animals (M1: Kruskal-Wallis test, H(2) = 504.29, p < 0.0001; M2: Kruskal-Wallis test, H(2) = 300.84, p < 0.0001), with all pairwise comparisons significant after Bonferroni correction. For M1, Toward vs Away p < 0.0001, Toward vs Neutral p < 0.0001, Neutral vs Away p < 0.0001. For M2, Toward vs Away p < 0.0001, Toward vs Neutral p < 0.0001, Neutral vs Away p < 0.0001. Overall, this was a very robust effect.

#### GAZE-HAND CORRELATIONS

If the gaze and hand system share a similar ego-allocentric weighting mechanism, their AWs should co-vary on a trial-by-trial basis. To examine whether the allocentric weighting of gaze and reach were coupled across trials, we correlated the gaze AW and reach AW across landmark shift trials for each animal. For M1, the overall correlation was weak but significant (Spearman’s ρ(2939) = 0.1083, p < 0.0001). For M2, the correlation was moderate and significant (Spearman’s ρ(2313) = 0.4504, p < 0.0001), suggesting that on trials where the gaze response was more strongly influenced by the landmark shift, the reach response was similarly influenced.

## DISCUSSION

The main purpose of this study was to analyze the influence of an allocentric cue, and a surreptitious shift of this cue on eye-hand coordination in monkeys. In summary, visual landmarks reduced gaze onset times, increased time to touch in both animals, and decreased temporal coupling of gaze and the hand. The presence of a stable landmark tended to reduce variable error (significantly only for M2 gaze), but had less influence on constant error, whereas landmark shifts negated this effect. In M2, but not M1, the stable landmark also increased the correlation of gaze and touch with target position. Gaze-to-hand touch correlations in landmark-centered coordinates were substantially reduced compared to spatial coordinates, particularly for M1, though most remained statistically significant. The temporal coupling between gaze and hand onset was strongest in the absence of a visual landmark and weakened significantly when a landmark was present. Both gaze and touch showed a significant shift in the direction of landmark shift in both animals. These shifts were only influenced by landmark distance in M2, but the touch shift was influenced by the direction of landmark shift relative to gaze in both monkeys, with higher weights when the landmark shifted toward the gaze position compared to away. Finally, the trial-to-trial allocentric weights of gaze and reach were modestly but significantly correlated. Each of these points is discussed in detail below.

### Memory-Guided Reaching in the Absence of Visual Landmarks

In our no-landmark task, the animal was only provided with a target (which turned off well before the gaze and reach movements) and a visual fixation point (which turned off just before the reach). This condition allowed us with a control for studying landmark influence but also provides some basic insights into eye-hand coordination.

#### TIMING

Previous experiments have found that a memory delay can either decrease or increase times compared to reactive visually guided movements (Krigolson & Heath, 2004; Lu & Fiehler, 2020). Our mean reaction times for memory-guided gaze (133.46 ms in M1; 136.99 ms in M2) and hand movements (204.12 ms in M1; 272.50 in M2) were comparable to or lower than typical reaction times to a visual stimulus (gaze: 107-166 ms; hand: 312-316 ms; Arora et al. 2019). This makes sense if reactive movements must fully process visual, cognitive, and motor information in a short time, whereas a memory delay allows one to develop a plan before it is triggered by the goal signal. Further, we found that longer delay times (from target onset to go signal) resulted in shorter reaction times in both gaze and hand movements. This is consistent with previous work showing that reaction time declines with increasing delay duration in both humans and non-human primates, suggesting that more lead time allows more complete motor preparation before the go signal (Crammond & Kalaska, 2000; Riehle & Requin, 1989; Rosenbaum, 1980). More lead time allowed animals to more fully formulate a coordinated motor plan before it is triggered by the go signal.

Eye-hand coordination usually involves getting gaze on target before the reach is complete, presumably to optimize foveal vision for the reach (Arora et al., 2019; Brouwer & Knill, 2007; Neggers & Bekkering, 2000; Song & McPeek, 2009). Foveal vision was not a factor in our memory-guided study, but gaze mean onset times still preceded hand onset by an average of 70.65 ms (M1) and 135.51 ms (M2). This difference falls within the wide, task related range (19-136 ms) reported in earlier human reaching studies (Angel et al., 1970; Prablanc et al., 1979; Rand & Stelmach, 2010) and is lower than what we found in a previous task where monkeys reached without a memory delay (Arora et al., 2019). It is noteworthy that some of this difference can be attributed to the long lead time between muscle activity and arm movement compared to the eye (Biguer et al., 1982), but these timing relationships can be altered or even reversed, depending on task complexity (Brouwer and Knill, 2007).

#### PRECISION AND ACCURACY

Note that in the current study, there was no visible target during the reach and saccade accuracy was not rewarded, so the only incentive for the animals was to reach accurately for a reward. Despite this, our animals both looked and reached toward the remembered targets, in the sense that gaze and touch showed significant correlations with target position. However, both gaze and hand landing points were less precise (larger ellipse fits) than those we reported in monkeys who reached directly to a sustained target (Arora et al. 2019). This agrees with previous memory delay studies in both saccades (Becker & Fuchs, 1969; Flanagan et al., 2008; Sharpe et al., 1975; Snyder et al., 2002) and reach (Chen et al., 2011; Obhi & Goodale, 2005; Vercher et al., 1995), especially in the absence of visual cues (Sheth & Shimojo, 2004; Flanagan et al. 2008). We did not observe increased errors with longer delays; perhaps the delays were not very long compared to human studies (Chen et al. 2011).

#### EYE-HAND COORDINATION

It is suggested that an internal sense of gaze direction is used to guide the reaching movement when we look at a target (Henriques et al., 1998; Henriques & Crawford, 2000; S. F. W. Neggers & Bekkering, 1999, 2000; Prablanc et al., 1979). In our memory-guided task, final gaze and hand position were significantly correlated across trials, suggesting a shared visual signal, at least at an early stage. A major source of this shared variance is likely the influence of the delay period on both gaze and hand reaction. They were also correlated in space, although it is possible that these were cross-correlations, since both showed higher correlations to target position. Also, as noted above, gaze variability was significantly higher than hand variability, so it is unlikely that gaze alone was guiding the hand movements. To gain further insights, we now return to our main question: how these various spatiotemporal factors are influenced by landmarks.

### Influence of a Stable landmark

#### TIMING

In the stable landmark condition, there was a significant decrease in gaze onset time (80.22 ± 115.33 M1; 109.96 ± 142.10 M2) compared to the no-landmark condition for both animals. This is near the range of ‘express saccades’ in monkeys (∼70 ms; Fischer & Boch, 1983), although the large variance suggests that some saccades were anticipatory, despite the variable delay time that preceded. In contrast, reach reaction onset was either unaffected overall (M1) or increased (M2). In general, these task and effector differences were preserved through the progressive drop in reaction time for longer delay periods, with the exception of M1’s reach onset times.

Chen et al. (2011) found reduced reaction times in the allocentric task compared to their egocentric task, especially in longer delays, whereas Obhi and Goodale (2005) did not find a significant effect of the landmark on their reach reaction time, and Krigolson and Heath (2004) also found that the hand movement duration was longer in the presence of a landmark. It may be that these task differences are due to differences in landmark information (present / absent during delays, small and near target vs. full field). In particular, a simple set of landmarks near the target facilitated saccade onset, whereas more complex scenes in other experiments might suppress movement through lateral inhibition. There may also be individual differences, as suggested in our reach data.

As a result of the reduced gaze reaction time and unchanged (or longer) reaction time for reach, landmarks increased the temporal gap between gaze onset and hand onset in both of our animals. The gap in M1 remained within the normal range reported in humans (Angel et al., 1970; Prablanc et al., 1979; Rand & Stelmach, 2010), while that in M2 exceeded it. As a result, saccades were more likely to be complete before reaches began. Further, reach end points were significantly delayed in the landmark conditions. Overall, this suggests that animals had more time before and during the reach to utilize vision for planning and feedback.

#### PRECISION AND ACCURACY

The presence of allocentric landmarks has been found to improve the accuracy and precision of reach and gaze responses (Byrne & Crawford, 2010; Krigolson et al., 2007; Krigolson & Heath, 2004; Li et al., 2017; Obhi & Goodale, 2005; I. Schütz et al., 2013). The stable landmark mostly increased accuracy, precision, and target correlations in this study, although this only reached significance in animal M2. Overall, these data are consistent with the notion that visual landmarks help stabilize goal representations during a memory delay (Chen et al., 2011; Li et al., 2017; Obhi & Goodale, 2005).

#### EYE-HAND COORDINATION

The spatiotemporal correlations between gaze and hand motion were significantly reduced in the stable landmark condition. Taken together with the temporal results (faster saccades, delayed reach) and spatial results (trends toward improved accuracy and precision) this suggests that animals developed a strategy that relied less on internal representations and more on visual feedback, which seems to make sense if the brain can code a remembered target relative to a visible landmark.

### Influence of the Landmark shift

#### TIMING

The stable landmark and landmark shift conditions were identical during target presentation and similar at the time of movement, the only difference being the reappearance of the landmark at a different location following the mask and before the final delay. It is likely for these reasons that gaze and hand timing were generally similar in these two tasks (with the exception that touch time was even longer for M2 in the Shift condition). So, we would again conclude that the timing of these events -whether by accident or design-optimized visual feedback during the reach.

#### ACCURACY AND PRECISION

In general, these were more similar (in the landmark shift task) to the no-landmark task than the stable landmark task, in both gaze and reach. From this one might conclude that the landmark shift reduced the reliability of the landmark, switching the system back to an egocentric state. However, a closer analysis showed that this was due to a more specific influence predicted from past literature, as follows.

#### ALLOCENTRIC WEIGHTING IN GAZE AND REACH

The increased aiming variance that we observed in the landmark shift task was not just random noise but rather reflected systematic shifts of gaze and reach in the direction of the landmark shift. To quantify this, we used the ‘allocentric weight’ (AW) score used in previous studies of this topic (Byrne & Crawford, 2010; Li et al., 2017). For reach, we found an overall mean AW 25.4% for M1 and 46.8% for M2, whereas the weights for gaze were 34.5% (M1) and 63.3% (M2). This is thought to represent the optimal weighting of allocentric, vs. egocentric information when these cues conflict (Byrne & Crawford, 2010; Li et al., 2017). Compared to a previous gaze study (Li et al., 2017), our reach AW data showed a similar dependence on landmark shift direction (toward or away) from gaze, but did not show a consistent dependence on landmark distance from gaze, perhaps suggesting subtle differences in mechanism.

Allocentric weighting is highly task-dependent, and is thought to increase with factors that increase egocentric noise, such as eye and body motion (Byrne & Crawford, 2010; Sheth & Shimojo, 2004) and increased memory delay (Byrne & Crawford, 2010; Chen et al., 2011; Fiehler et al., 2014; Lu & Fiehler, 2020). Allocentric weight also increases with the scale of large-scale and dynamic environments using virtual reality and naturalistic scenes (Karimpur et al., 2020; Lu & Fiehler, 2020). In this case, the shift of four salient dots around our target was designed to produce a strong weighting, and the higher gaze AW may relate to the higher level of egocentric noise we observed in gaze.

#### ALLOCENTRIC WEIGHT AND EYE-HAND COORDINATION

If the gaze and reach systems share a common visual mechanism for allocentric-egocentric integration, their trial-by-trial allocentric weights should be positively correlated. We found this to be the case for both animals, though the strength of the correlation differed substantially. The stronger AW coupling in M2 is consistent with its higher allocentric weights overall and suggests a greater degree of shared allocentric processing between the two systems, whereas the weaker coupling in M1 may reflect its lower overall AW, i.e., higher levels of independent egocentric noise. These results suggest that while gaze and reach AWs are not highly correlated, they share at least some component of this mechanism.

### Possible Neural Mechanisms

To speculate about the neural mechanisms for the behavior described above, it is useful to divide the effects into 1) general influence of landmark on eye-hand coordination, and 2) the specific mechanisms for egocentric-allocentric integration. In either case, we need to consider where landmark signals could interact with the eye-hand coordination system.

The topic of eye-hand coordination is itself controversial but is generally thought to involve some interaction of gaze and hand signals in premotor frontal areas, posterior parietal cortex, and the cerebellum (Miall et al., 2001; Pesaran et al., 2006; Snyder et al., 2002). Human studies suggest that allocentric visual codes originate in the occipito-temporal ‘ventral stream’ (Chen et al., 2014; Chen & Crawford, 2017; Goodale & Milner, 1992; Schenk, 2006) and converge with egocentric visuomotor systems in parietal and frontal cortex (Chen et al., 2018; Musa et al., 2024, 2025). Consistent with this, Taghizadeh et al. (2024) found evidence of object-centered coding in reach areas of parietal and premotor cortex. Finally, a gaze paradigm very similar to the one used here revealed integration of target and landmark signals in the prefrontal visual responses (A. Schütz et al., 2023), followed by responses to shifting landmarks in the memory delay and reintegration in the motor response (Bharmauria et al., 2020, 2021).

The gaze and reach timing changes that we observed in the presence of a landmark have the appearance of a deliberate motor strategy that could result from our training regime or a lifetime of experience, and involve context dependent signals through cerebro-cerebellar systems (Tanaka et al., 2020; Wolpert et al., 1998). Likewise, the modest decrease in spatial eye-hand coordination observed in our stable landmark task could be due to decreased reliance on the integration of egocentric signals and increased reliance on the use of landmark signals. The latter influence was only partially correlated between gaze and hand, perhaps because it relies on early visual mechanisms (named above) feeding into separate parietal and frontal motor centers. However, we cannot dismiss that these correlations arose from lateral gaze-hand connections in the areas mentioned above. Finally, although ego-allocentric integration involves specific cellular and population mechanisms (Bharmauria et al., 2020, 2021; A. Schütz et al., 2023), properties arise when deep neural nets are trained to perform this task (Abedi Khoozani et al., 2022), suggesting a general solution that likely extends to the reach system (Bharmauria et al., 2025).

## CONCLUSIONS

The tasks that we employed in this study demonstrated the expected stabilizing influence of stationary landmarks and destabilizing influence of shifting landmarks on both gaze and hand behavior. Further, the latter influence was at least partially shared between gaze and hand, suggesting at least a partially shared mechanism and a similar algorithm, consistent with previous studies of ego-allocentric integration. Less expected was the influence of visual landmarks on the motor aspects of eye-hand coordination, including changes in timing and spatial correlations, which might relate to specific sensorimotor strategies to optimize available information. Based on past literature, one expects these observations to translate well to humans, making this non-human primate paradigm a promising experimental model to investigate neural mechanisms.

## ACKNOWLEDGEMENTS

The authors thank Dr. Vishal Bharmauria for advice.

## FUNDING STATEMENT

This project was supported by VISTA and the Connected Minds Program, which were supported in part by the Canada First Research Excellence Fund. J.D. Crawford was supported by a York Research Chair.

## Notes

### Competing Interest Statement

The authors have declared no competing interest.

