## Supplemental Figures 1-8 for "Visual Landmarks Influence Eye-Hand Coordination in Memory-Guided Reaches"

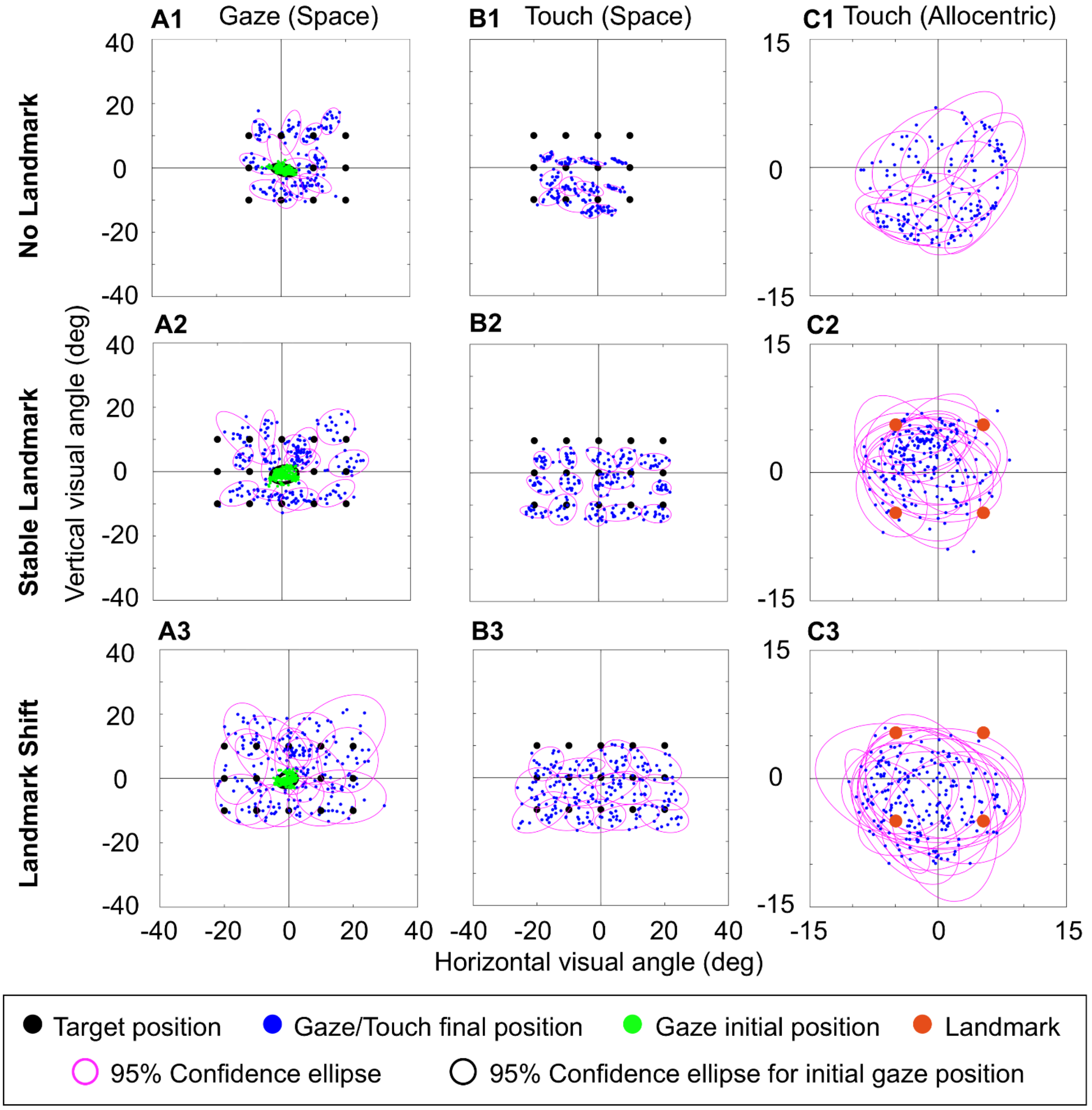


**Supplemental Figure 1**. Example 95% confidence ellipses for data collected in one session in M2. (A) gaze endpoints at the end of saccades in each condition (B) touch endpoints at the point of screen contact. (C) touch endpoints at the point of screen contact in allocentric coordinates.

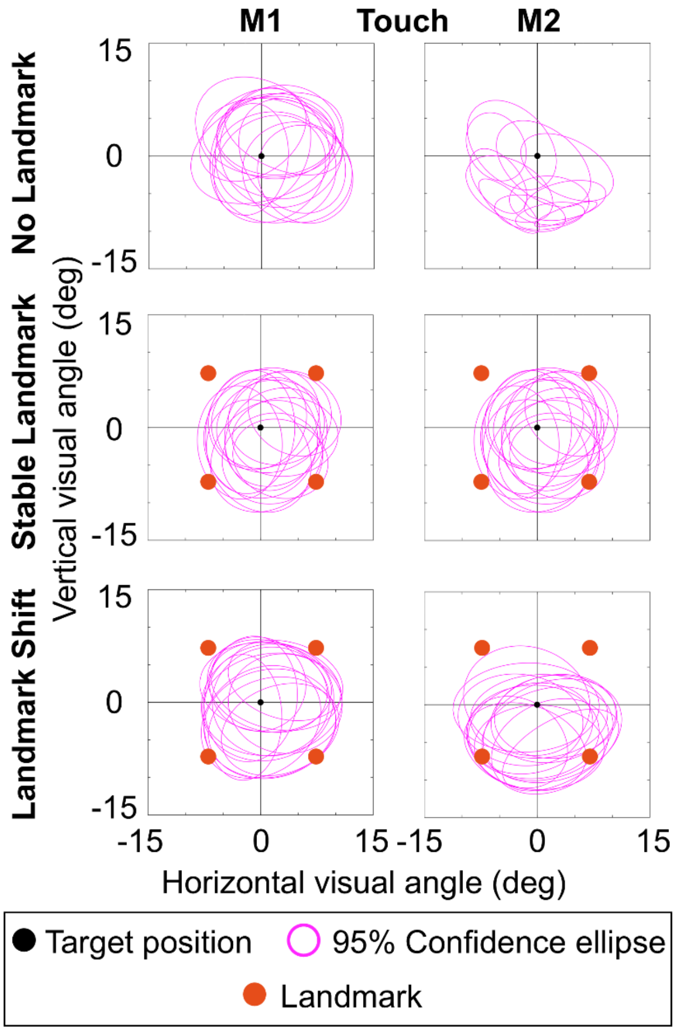


**Supplemental Figure 2**. 95% confidence ellipses showing the touch endpoints at the point of screen contact in allocentric coordinates for the complete data set of M1 and M2.

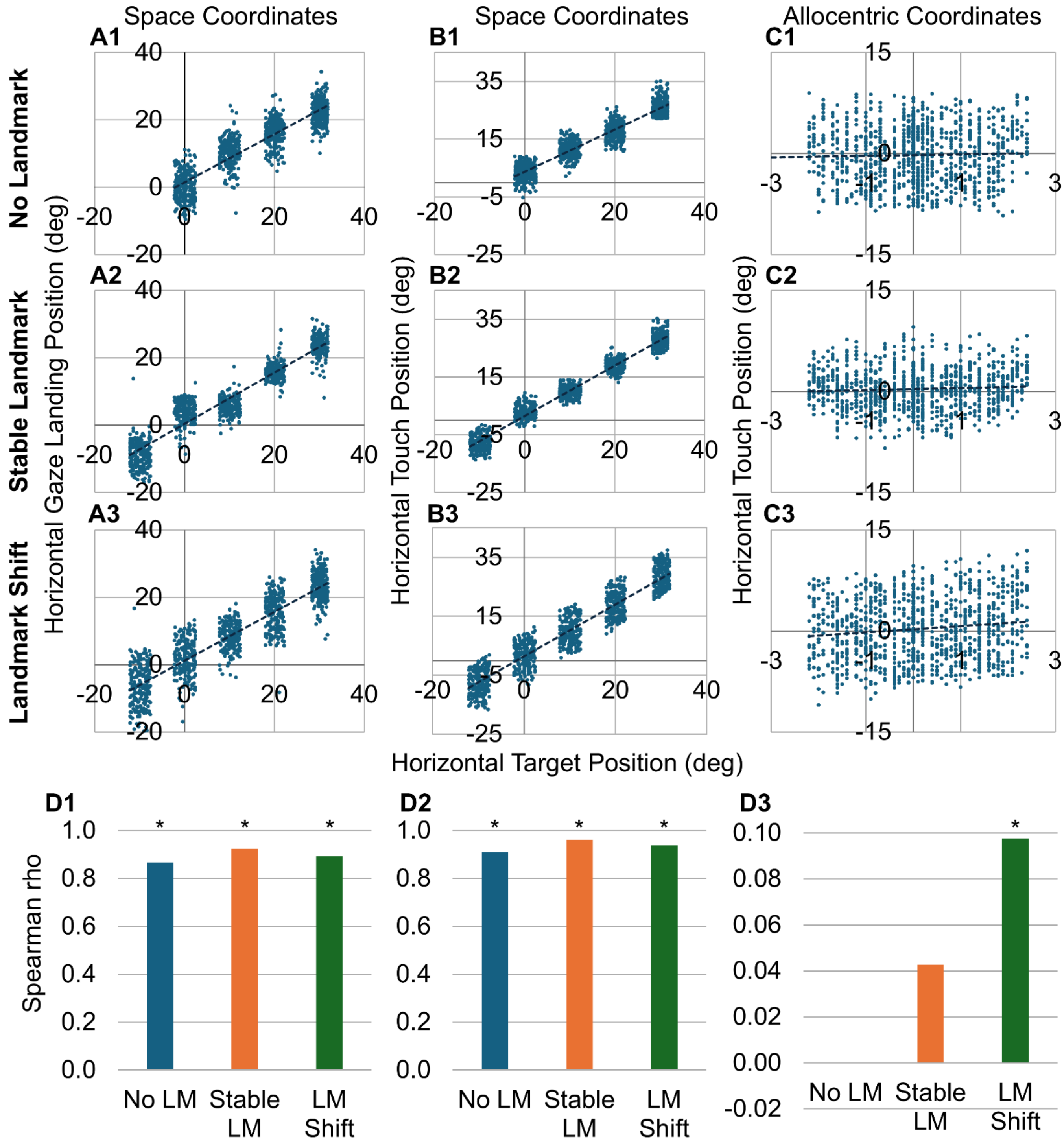


**Supplemental Figure 3**. Correlation of hand and gaze position to target position in M2. Shown in the horizontal dimension. (A) Correlation of gaze landing to target position in space. (B) Correlation of hand touch to target position in space. (C) Correlation of hand touch to target position in landmark coordinates. (D) Bar graphs summarizing the Spearman correlation for each condition. Asterisks indicate significant correlations ($p < 0.05$).

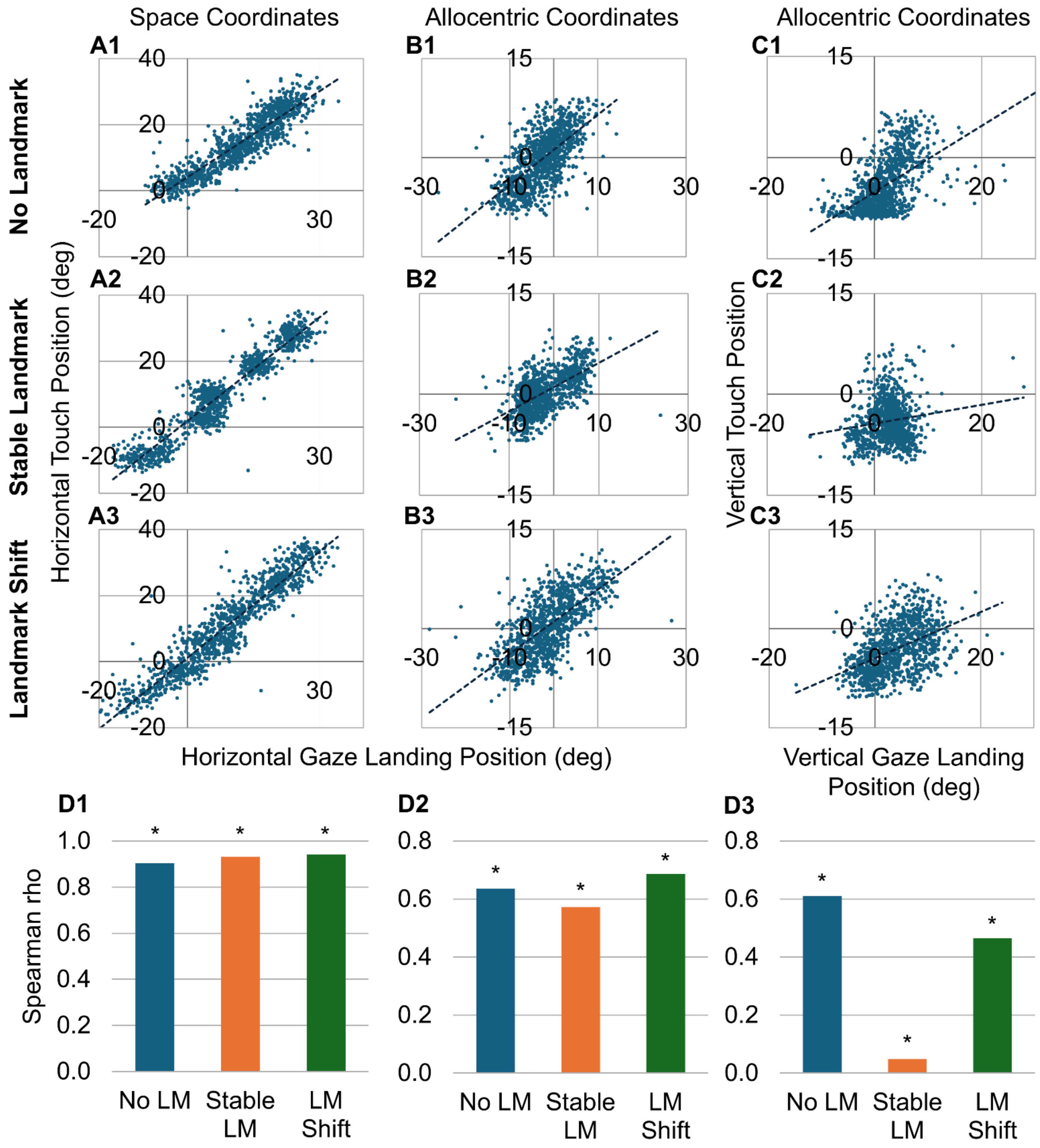


**Supplemental Figure 4**. Correlation of hand touch position to gaze landing position in M2. (A) Correlation of horizontal hand touch to gaze position in space. (B) Correlation of horizontal hand touch to gaze position in allocentric (landmark-centered) coordinates. (C) Correlation of vertical hand touch to gaze position in allocentric coordinates. (D) Bar graphs summarizing the Spearman correlation for each condition. Asterisks indicate significant correlations ($p < 0.05$).

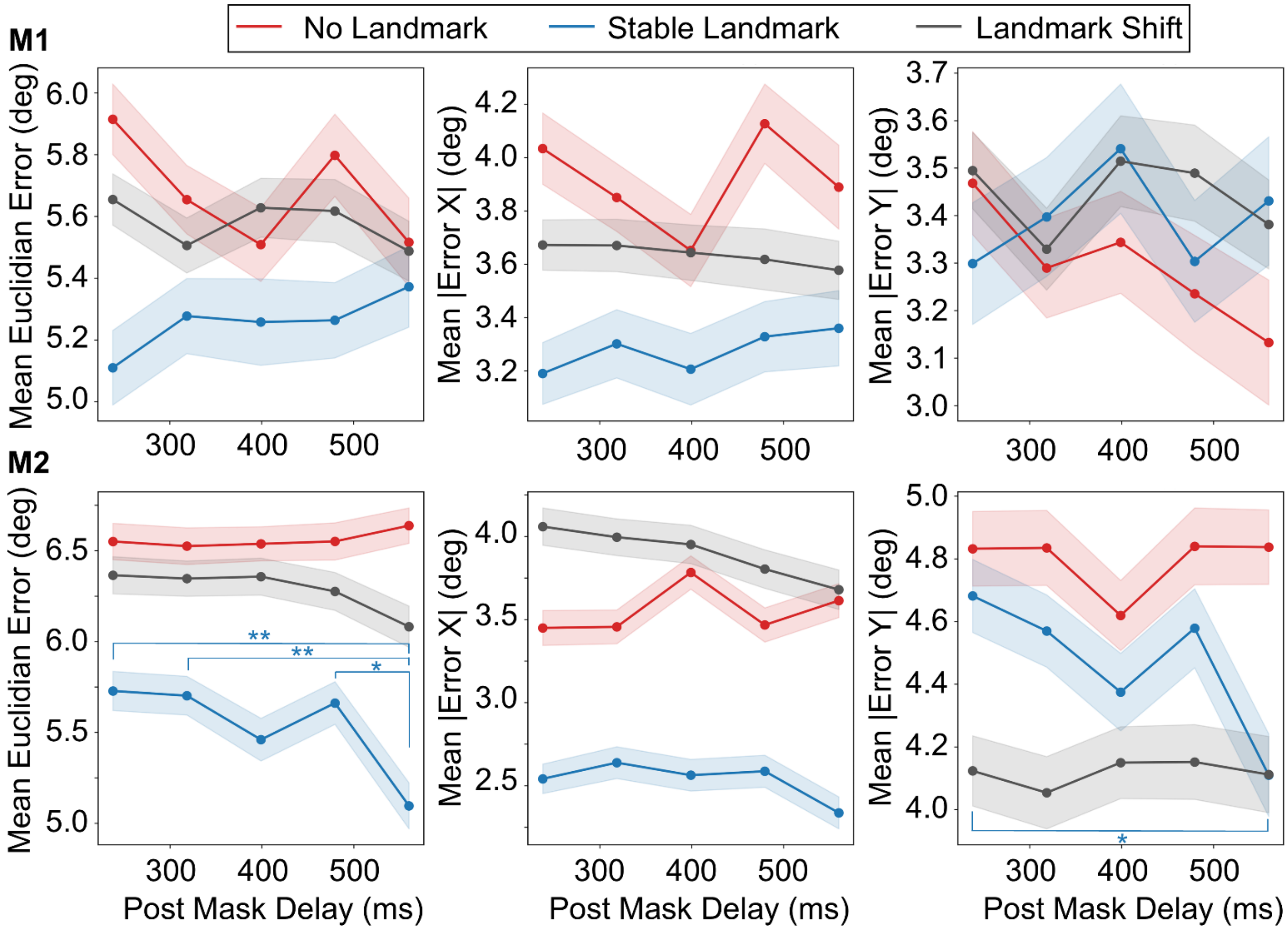


**Supplemental Figure 5**. Mean reach error as a function of post-mask delay for M1 and M2. Each panel plots error against the five delay bins (198–278, 279–359, 360–439, 440–520, and 521–600 ms) for the no-landmark (red), stable landmark (blue), and landmark shift (grey) conditions. Columns show, from left to right, mean Euclidean error, mean horizontal error (|Error X|), and mean vertical error (|Error Y|), all in degrees. Points show the bin means and shaded regions indicate ± SEM. Asterisks mark significant differences between post-mask delay bins in the stable landmark condition (Dunn's post-hoc test, Bonferroni-corrected; * $p < 0.05$, ** $p < 0.01$). All other comparisons, $p > 0.05$.

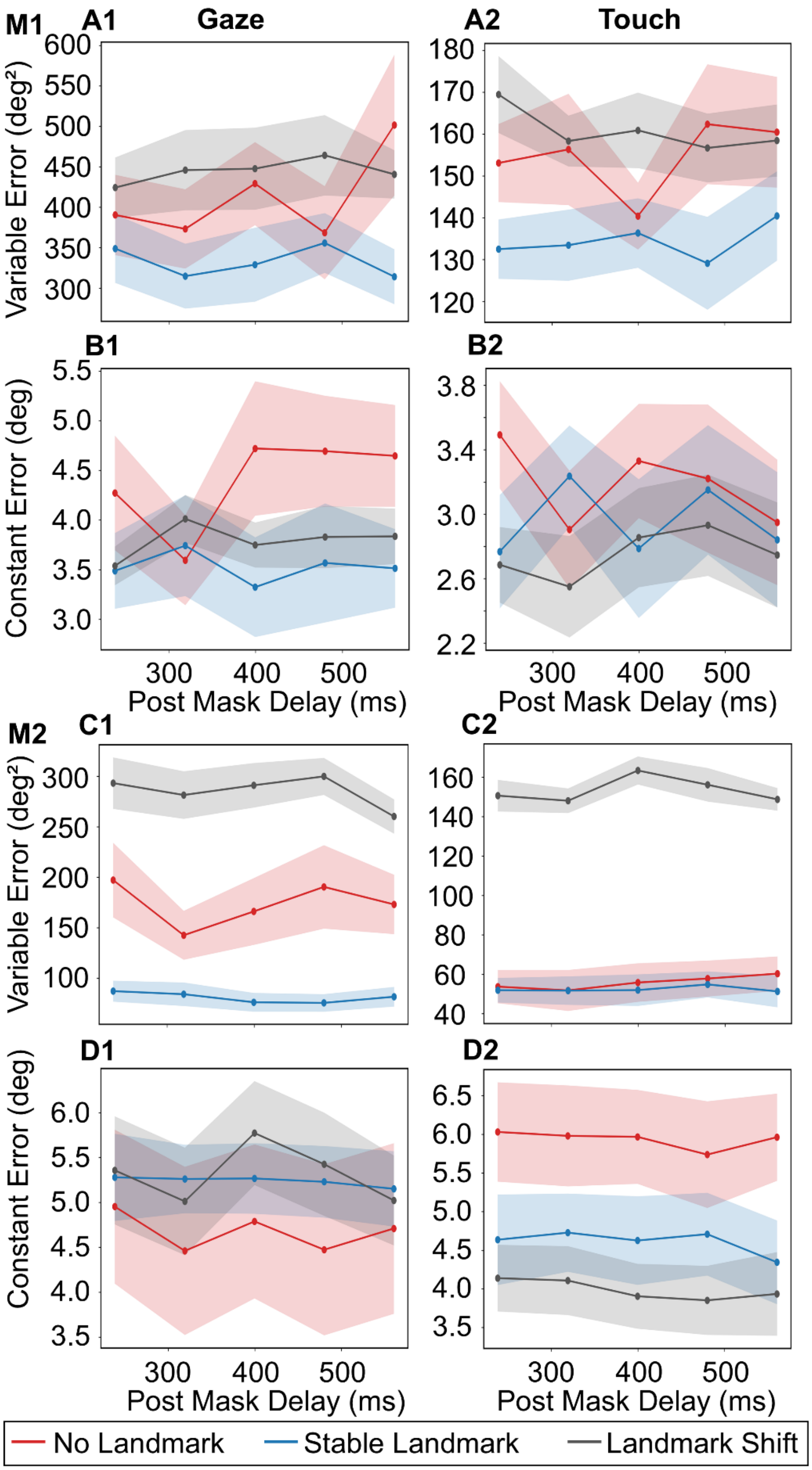

M1
A1
Gaze
A2
Touch
Variable Error (deg²)
B1
B2
Constant Error (deg)
Post Mask Delay (ms)
M2
C1
C2
Variable Error (deg²)
D1
D2
Constant Error (deg)
Post Mask Delay (ms)
No Landmark
Stable Landmark
Landmark Shift

**Supplemental Figure 6**. Variable error (ellipse area, A and C) and constant error (distance from ellipse center to target, B and D) as a function of post-mask delay duration for M1 (A and B) and M2 (C and D) separated by effector (Gaze, 1; Touch, 2). Trials were divided into five delay bins (198–278, 279–359, 360–439, 440–520, and 521–600 ms), and mean variable and constant error were computed within each bin for the no-landmark (red), stable landmark (blue), and landmark shift (grey) conditions. Points show bin means and shaded regions indicate ± SEM.

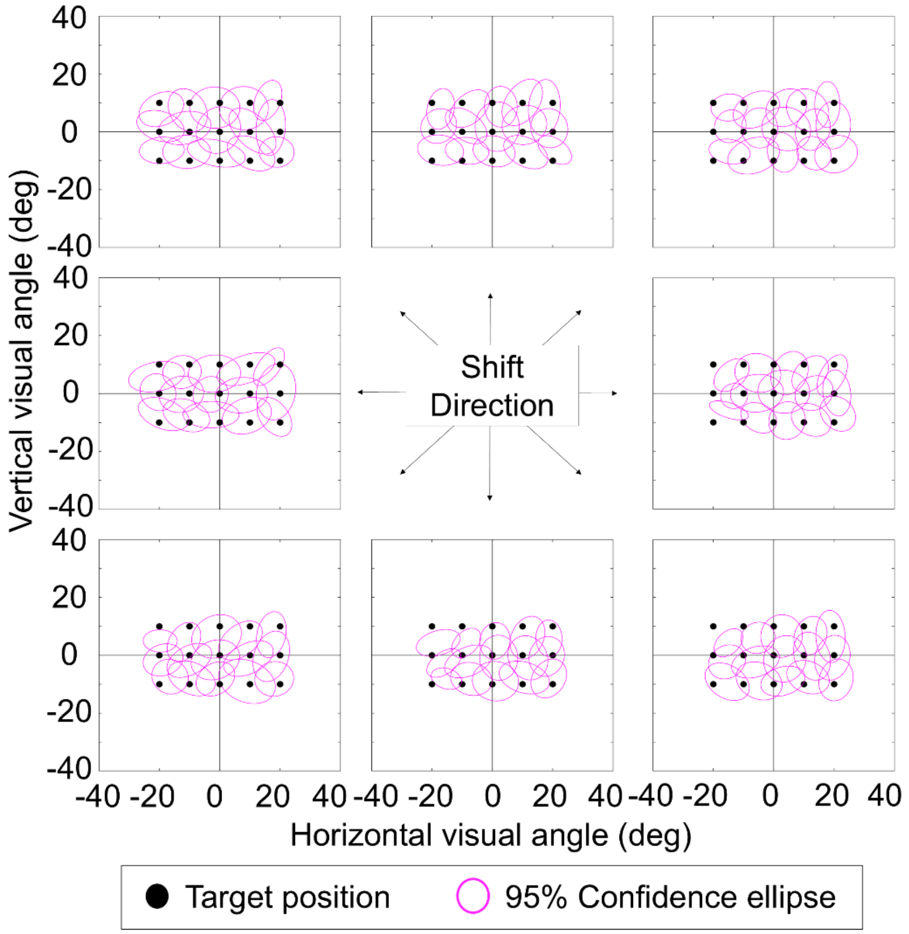


**Supplemental Figure 7**. Influence of different landmark shift directions on 95% confidence ellipses for touch. Data from the landmark shift condition collected from M1 were separated into the eight shift directions.

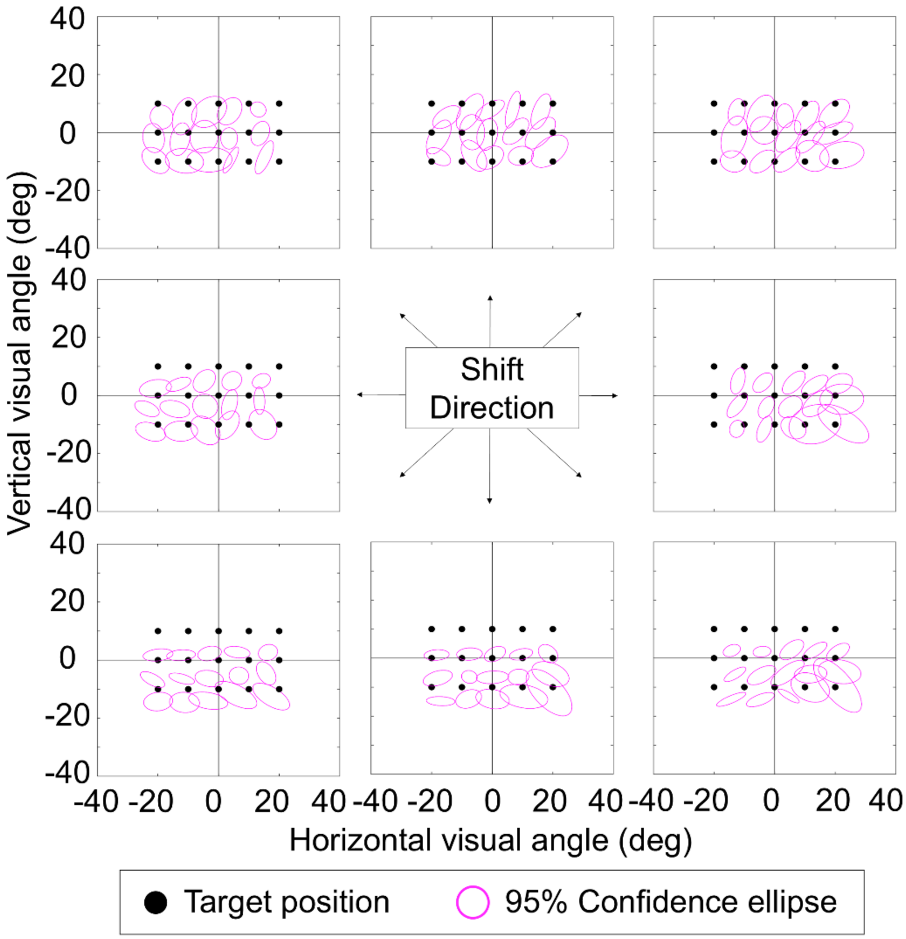


**Supplemental Figure 8**. Influence of different landmark shift directions on 95% confidence ellipses for touch. Data from the landmark shift condition collected from M2 were separated into the eight shift directions.
